# Differential fate of acellular vascular scaffolds in vivo involves macrophages and T-cell subsets

**DOI:** 10.1101/2020.11.21.392654

**Authors:** Debashish Banerjee, Nikhil B. Nayakawde, Deepti Antony, Meghshree Deshmukh, Sudip Ghosh, Carina Sihlbom, Evelin Berger, Uzair Ul Haq, Michael Olausson

## Abstract

Biological scaffold or implant is a popular choice for the preparation of tissue-engineered organs and has the potential to address donor shortage in the clinics. However, biological scaffolds prepared by physical or chemical agents cause damage to the extracellular matrix by potentially inducing immune responses after implantation. The current study explores an alternative route for the preparation of acellular scaffolds and explores the fate of the prepared scaffolds in a milieu of immune cells following implantation without using immunosuppressant. Using the syngeneic (Lewis male-Lewis female) and allogeneic (Brown Norway male-Lewis female) models and different tissue routes (subcutaneous vs omentum) for transplantation, normal blood vascular scaffolds were implanted which was converted to acellular vascular scaffolds by *in vivo* natural decellularization at the end of 2 months of observation. We also prepared chemically decellularized acellular scaffolds from normal untreated blood vascular scaffolds using a cocktail of chemicals which was also similarly placed in subcutaneous and omentum sites. Here, we applied in-depth quantitative proteomics along with histology and image analysis to comprehensively describe and compare the proteome of the natural and chemically decellularized scaffold. Our data confirm that site-specific advantages exist in modulating the ECM and regulating the immune responses (macrophage and T cells) following implantation, which possibly led to the production of an acellular scaffold (natural decellularization) under *in vivo* conditions. The current approach opens up the possibility to create tailor-made acellular scaffolds to build functional blood vessels. In addition, the identification of different tissue sites facilitates differential immune response against the scaffolds. This study provides a rich resource aimed toward an enhanced mechanistic understanding to study immune responses under similar settings in the field of transplantation and regenerative medicine.

**Impact statement:** The development of a scaffold helps in the preparation of a functional organ in the clinics. In the current study, we prepared an acellular vascular scaffold by utilizing site specific tissue changes and vis-à-vis compared with a conventionally chemically prepared biological scaffold at genomic and protein level, which helped us to identify immunological trigger following implantation. The current study which was carried out without any immunosuppressive agents could help to establish (a) alternative strategies for preparing biological scaffolds as well as (b) implantable sites as potential bioreactors to circumvent any adverse immune reactions for acceptance of the scaffold/implant post implantation.

## Introduction

Synthetic grafts and native blood vessels are often used as a conduit for revascularization as a regenerative measure or tissue engineering approach to create vascular scaffold identical. The initial development of blood vascular scaffold^1^ and refinement in terms of rapid endothelization and extracellular matrix (ECM) deposition^2^ has helped tissue engineering technique to develop successful blood vessels with minimum thrombosis, inflammation, and scaffold acceptance.^3, 4^ Despite the inherent drawbacks, biological scaffold prepared by decellularization technology has been successfully used to recreate various types of tissues and organs, with an approach to repopulate with a patient’s cells to produce a personalized vascular scaffold.^4, 5^

The usual chemical procedure used for the development of acellular biological scaffold (decellularization technology) have been reported to damage the extracellular matrix^6^ and elicit immunological reaction and incompatibility with limited mechanical tolerance^7, 8^. However, natural vascular scaffolds in comparison to synthetic scaffolds have certain advantages such as natural binding sites for cell adhesion (conserved native 3-D structure), improved biomimetic and biocompatibility stimulating colonization, promoting the proliferation of recruited cells. Remodelling of autologous vascular scaffolds *in vivo* which acquired native tissue architecture with maintenance of smooth muscle cells, extracellular matrix organization, and endothelial cells have been reported.^4, 9^

Selection of scaffold implantation sites with unique cellular niche and different blood supply e.g., subcutaneous and omentum has been reported to support variety of free structures for reconstructive purposes^10–12^. The uniqueness of the two sites for implantation has earlier been documented with the omentum richer in blood supply^13^ whereas the subcutaneous site associated with low oxygen tensions.^14^

Implantation of a biomaterial and its degraded by-product induces an immune reaction in the host which determines the final integration and the biological performance of the implant.^15, 16^ The nature of the host immune cells e.g. macrophages depends on the biomaterials influencing the outcome of the tissue-specific innate and adaptive immune responses.^17^ A modern scientific tool to assess biological scaffold includes mass spectrometric based quantitative proteomic analysis and bioinformatics prediction to identify and analyze proteins of the extracellular matrix^18^ as well as an immune response after transplantation.^19^

During the study, we developed approaches to prepare scaffolds and study the fate of scaffolds following implantation by proteomics and immunohistological tools to monitor expression of macrophage and immune cell markers. The findings from our study, could help us to better understand the remodeling of the implant because of the putative interaction between the ECM and the immune infiltrating cells, chose an alternative source of implantation, as well as manipulate the immunological target to minimize adverse immune reactions in the clinics.

## Materials and Methods

All reagents used were of analytical grade. Antibodies used are summarized in **Supplementary Table S1.**

All animal work followed the accepted guidelines reviewed by the **local animal welfare committee at the University of Gothenburg** (Göteborgs Djurförsöksetiska Nämnd, Ethical Number, 151/14). The donor male (Lewis and Brown Norway) rats and recipient female Lewis rats were operated under general anaesthesia with inhalant anaesthetic, isoflurane. Details of animal handling and operation is included in the **Supplementary file.**

### Chemical decellularization procedure of donor aorta

The DC steps were performed as reported earlier^20^ at room temperature (20-24°C) with slight modifications using a closed-circuit system to minimize contamination. Each decellularization cycle consisted of a 7-hour treatment with 0.2% SDC (Sigma-Aldrich, St. Louis, MO, USA) with an alternate cycle of 7-8-hour treatment with glycerol mix solution (10% glycerol, 0.9% sodium chloride and 2 mM sodium-ethylene diaminetriacetic acid (Sigma-Aldrich, St. Louis, MO, USA) with a wash with distilled water in between to remove cell debris as shown schematically in **Fig. 1**. After the DC cycle, washing was performed overnight with sterile water using an open system to drain out the wash continuously. Subsequently, the chemically decellularized aortas were treated with 900 units (25 μg/mL DNase I recombinant, RNase free, Cat. No. 4716728001, Roche, Merck KGaA, Darmstadt, Germany) using a total of 3 DNase treatment for 2 hours each to achieve the desired level of DNA assessed by (Quant-iT™ PicoGreen™ dsDNA Assay Kit, Catalog number: P7589) and H&E staining. A total of 5 cycles was given to achieve decellularization as assessed from DNA load and H&E staining. The DC and normal (untreated scaffolds) were subsequently sterilized before implantation as outlined in **Supplementary file**.

**Figure 1.**
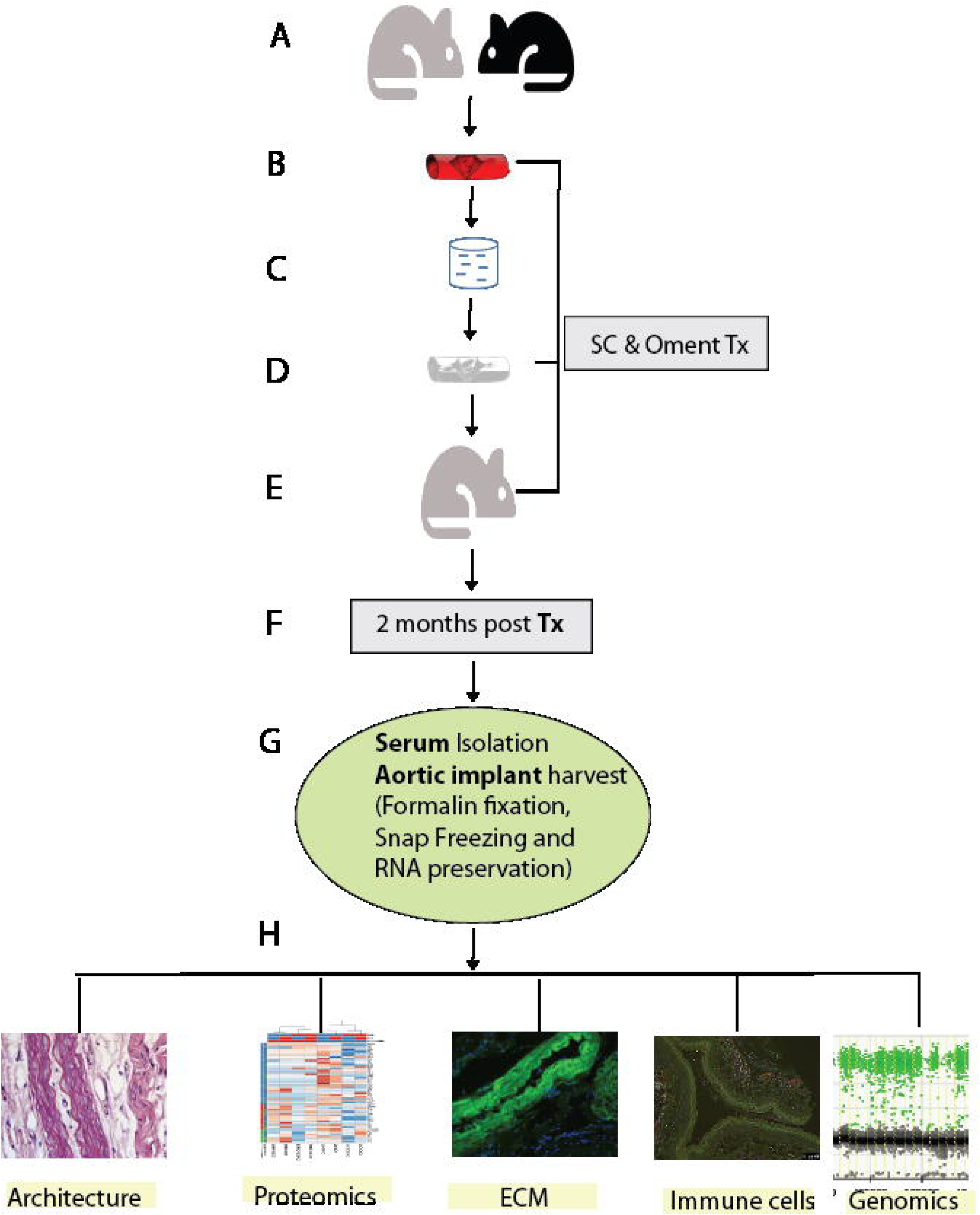
The experimental design and layout of the study. The experimental design is presented in a sequential layout **(A-H)**. Male Lewis and Brown Norway donor rats **(A)** were used for harvesting the aorta **(B)**. The aortasfrom donor rats (male Lewis/Brown Norway) were then perfused with a chemical solution **(C)** to prepare a decellularized aortic scaffold **(D).** Implantations of the donor aorta **(B)** and decellularized aortas **(D)** at the subcutaneous and omentum route were performed in female recipient mice **(E).** The implanted aortas and serum were collected following sacrifice of the recipient mice after 2 months, (**F)** for downstream processing for serum and tissues, **(G).** The tissue architecture, proteomics, ECM, and genomic characterization were performed, (**H).**

### Implantation and post-implantation biochemical study

The Lewis female recipient rats (n=6) were used for the implantation study as outlined in the Supplementary file.

The nomenclature of the donor as well as implanted scaffolds is outlined below and in **Supplementary Table S2**. The implants are named according to the genetic origin (syngeneic and allogeneic), nature (N or DC) of scaffolds and sites of implantations (subcutaneous, SC or Omentum, O). The Lewis (donor) to Lewis (recipient) is designated as syngeneic whereas Brown Norway (donor) to Lewis recipient is designated as allogeneic. The syngeneic implants at the (a) SC sites - LNSC and LDCSC; (b) O site - LNO and LDCO; allogeneic implants at the (a) SC sites - BNNSC and BNDCSC; (b) O site - BNNO and BNDCO.

### Processing of samples

In addition to animal groups mentioned above, we also prepared a sham treatment group consisting of female rats of same age as the recipient animals. The control, sham and implanted scaffolds (≈3-4 mm) each were stored separately for (i) proteomics and DNA snap-frozen in liquid nitrogen and stored at -150°C; (ii) for gene expression were stored with RNALater at - 80°C. For histological analysis (≈6 mm) were immediately fixed in formalin. The serum samples were prepared from collected blood and stored at -20°C for further analysis.

### Histological assessment

#### Tissue processing and staining

The control scaffolds and implants (collected from SC and omentum sites from the recipient mice) were fixed in formalin for a period of 48 h before paraffin embedding and sectioning at 4 μM thickness. The sections were finally processed for H&E staining and immunohistochemical localization of different protein targets (ECM and immune cells) as outlined in **Supplementary file**.

### Architectural preservation and nuclear basophilia score of the decellularized aorta

The H&E stained sections were evaluated for structural integrity (architectural preservation) and nuclear loss (nuclear basophilia) blindly by a pathologist (A.K.) using a Comprehensive method of masking tissues^21^ with individually labelled slides followed by Ordinal Data Measurements.^22^

The semi-quantitative evaluation of the tissue architecture was performed using a scoring represented as (1) complete breakdown of tissue, (2) marked disruption, 3) moderate disruption, (4) minimal disruption, and (5) no damage to cellular architecture.

Similarly, a semi-quantitative nuclear loss score was performed representing nuclear basophilia: 1- complete loss (100%), 2- substantial loss (70%), 3-moderate loss (50%), 4- minimal loss (30%), and 5- no loss of nuclear basophilia (0%).^22^

### Inflammatory, fibrosis and necrosis score

To monitor the degree of inflammation, fibrosis and necrosis, a simple scoring system was used based on the following scoring criterion- (i) absent/ none-0, (ii) mild/weak- 1, (iii) moderate- 2 and (iv) severe/strong- 3. Qualitative assessment for the type of inflammatory cells was also monitored.

### Immunohistochemical assessment

Formalin-fixed paraffin sections (4 μM) were deparaffinized and target protein was probed by primary and secondary antibody before being subjected to image acquisition and analysis as outlined in **Supplementary file.**

### Proteomic Analysis

For proteome analysis, 3-4 mm of donor scaffolds and implants were used. In total, twelve groups were selected for proteomic analysis **(Supplementary Table S2)**. Samples were digested with the filter-aided sample preparation (FASP) method^23^ and peptides were labeled using TMT 11-plex isobaric reagents (Thermo Fisher Scientific) according to the manufacturer’s instructions. MS analyses were performed on an QExactive HF mass spectrometer equipped with Easy-nLC 2000 (Thermo Fisher Scientific) using a gradient from 7% to 35% B over 76 min followed by an increase to 100% B for 8 min at a flow of 300 nL/min followed by MS2 data dependent acquisition method. Identification and relative quantification were performed using Proteome Discoverer™ version 2.2 (Thermo Fisher Scientific). Details outlined in **Supplementary file.**

### Gene expression analysis - calculation of target gene

#### RNA Quantification and ddPCR

Tissue samples (scaffolds/implants) stored at -80°C in RNAlater™ Stabilization Solution (Catalog number: AM7021, Thermo Fisher Scientific, 168, MA USA 02451) were used for analysis RNA isolation and quantification by ddPCR. The calculated input RNA (ng) was calculated initially from the theoretical input RNA(ng) and reference, β-actin copy number as elaborated.^24^ Details outlined in the **Supplementary file.**

### XY chromosome quantification-identifications of the resident cells around the implant

The infiltrating cells in the implants were identified for their origin i.e. either of donor (male) or recipient (female) and expressed as percentage copies of the Y chromosome by ddPCR. Details of XY chromosome primers and amplification outlined in **Supplementary file.**

### Proteome Profiler of cytokines present in serum

Cytokine arrays were used to find semi-quantitative data of cytokines and chemokines in serum as an index of the immune response following graft implantation and compared with sham treatment. Details outlined in **Supplementary file.**

### Statistical analysis

The data were statistically analyzed using GraphPad Prism v.8.4 (GraphPad Software, La Jolla, CA) and R statistical language. Differences between two groups were analyzed and p-values below 0.05 were considered statistically significant. The statistical analysis carried out in each experiment is defined in the corresponding figure legends. Details outlined in **Supplementary file.**

## Results

The overall experimental set-up is shown **(Fig. 1)** with nomenclature of the different scaffolds and implants outlined in **Supplementary Table S2**.

### Decellularization strategy-Modulation of scaffolds following implantation

Chemically decellularized scaffolds (DLDC and DBNDC) from normal control (DLN and DBNN) aorta/ scaffold were prepared with the DNA content found to be below 10 ng which was significantly lesser (syngeneic - ^⍰^p<0.05 and allogeneic-^⍰⍰^p<0.01) than the corresponding normal control (N) scaffolds **(Supplementary Fig. S1).**

Semi-quantitative histological scoring also confirmed (79.31%, p^⍰⍰^<0.01) of the acellularity (nuclear basophilia) of the DC control scaffolds compared to the N control scaffolds (**Fig. 2A and Fig. 2B, Supplementary Fig. S2**).

**Figure 2.**
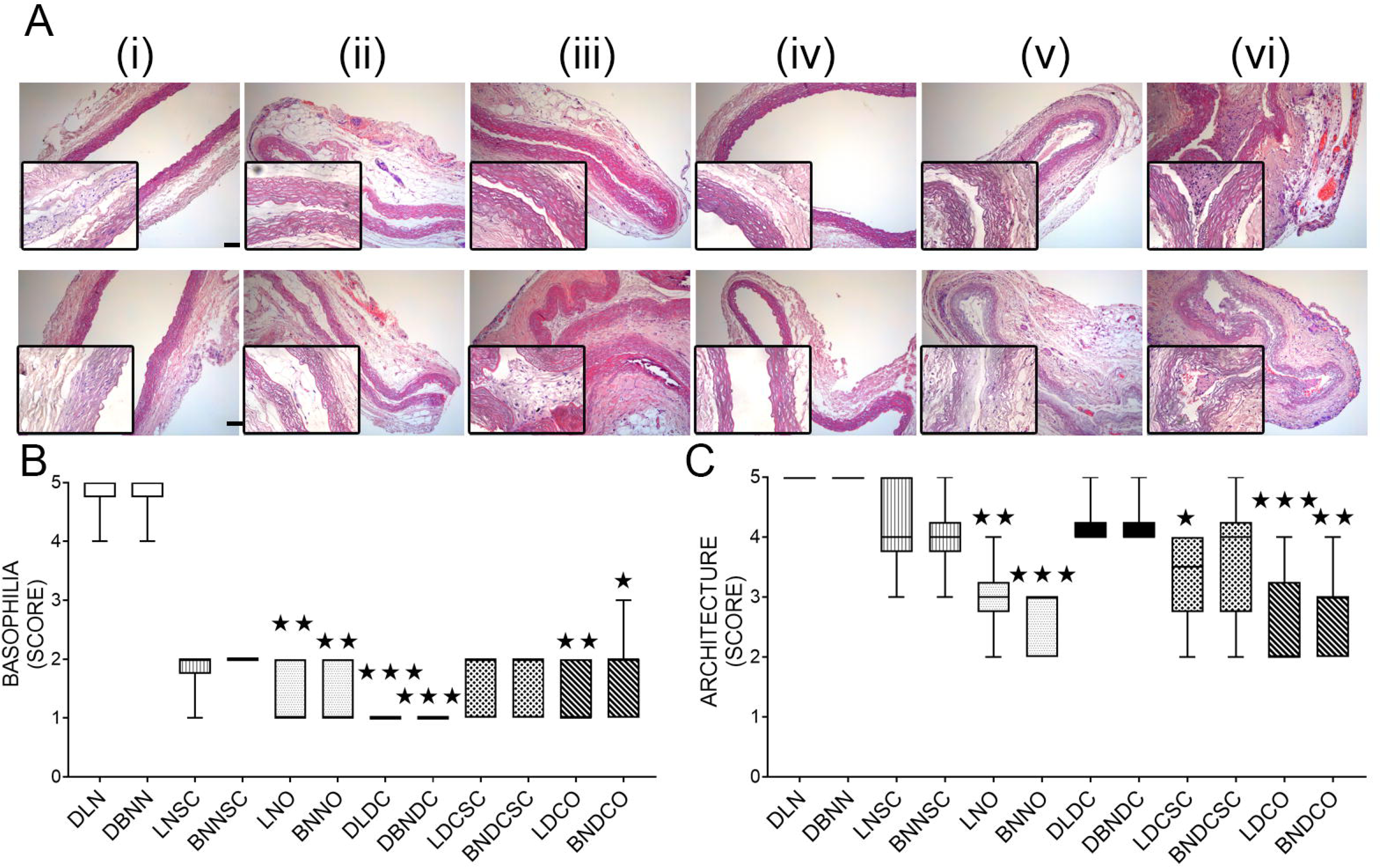
H&E staining of normal and decellularized control scaffolds and implants. (**A**). Upper panel-Lewis (L) and lower panel-Brown Norway (BN) normal anddecellularized scaffolds andafter implantation and harvest from subcutaneous and omentum route after 2 months. (i) normal (N), (ii) normal subcutaneous, (iii) normal omentum (NO), (iv) decellularized, (v) decellularized subcutaneous (DCSC) and (vi) decellularized omentum (DCO). All the donor scaffolds were stained with H&E and compared to the implanted aortas in terms of the Nuclear basophilia (**B**) and Architectural score (**C**) as defined in the Materials and Methods section. Representative images are shown at 10× magnification. Scale bars: 100 µm. Results are presented (n=6in each group) as maximum and minimum of the rank. (median in the middle of each box). p<0.05, p<0.01 and p<0.001 indicates a significant difference in comparison to the respective normal donor scaffolds by Kruskal-Wallis test followed by Dunn’s Multiple Comparisons post hoc tests.

The tissue architecture score (**Fig. 2A and Fig. 2C, SupplementaryFig. S2**) of the control, DC scaffolds were found to be reduced compared to the control, N control scaffolds. Following implantation, the omentum (O) implants showed a reduction (syngeneic and allogeneic, N and syngeneic, DC - 72.41%, p<0.01; allogeneic, DC- 68.96, p<0.05) compared to the control, N controls (**Fig. 2A and 2B, Supplementary Fig. S2**).

The architecture of O implants for the syngeneic (N- 40%, p <0.01; DC- 50%, ^⍰⍰⍰^p<0.001) and allogeneic (N- 46.67%, ^⍰⍰⍰^p<0.001; DC- 43.33%, ^⍰⍰^p<0.01) was significantly lower compared to the N controls. The subcutaneous (SC) implants (N syngeneic, allogeneic and DC allogeneic) did not show any change in the tissue architecture except for DC syngeneic implant (33.33%, p<0.05) compared to the respective N controls.

The N allogeneic O implant and the DC syngeneic (except SC) and allogeneic implants showed higher level of inflammatory cells with evidence of a few giant cells, compared to the N controls, which were mainly lymphocytes **(Supplementary Fig. S3)** and later characterized immunohistologically to be macrophages.

The N allogeneic O implant showed a significant increase in the fibrosis score compared to the respective N control (4.25 folds, p<0.01) and N allogeneic SC implant (3.4 folds, p<0.05). The allogeneic O implants (N- 6 folds, p<0.01 and DC- 5.5 folds, p<0.05) and DC SC implants (5-5.5 folds, p<0.05) showed significantly more necrosis compared to the N controls. None of the SC implants showed significant necrosis except for the syngeneic DC subcutaneous implant which showed higher necrosis score (5 folds, p<0.05) compared to the normal SC implant (LNSC) (**Supplementary Fig. S3**).

To confirm the absence of donor cells (male), Y-chromosome presence was monitored by ddPCR analysis which revealed negligible donor cells suggestive of an efficient natural decellularization *in vivo* **(Supplementary Fig. S4).**

The sham treated rats which received identical surgical intervention as the recipient rats showed identical DNA content and tissue architecture of aorta compared to the normal control aorta (**Supplementary Fig. 5).**

### Proteomic analysis of syngeneic and allogeneic scaffolds following implantation

To study the differential expression of the protein after implantation, the controls and implants were subjected to 11-plex TMT-based LC-MS/MS-based proteomic analysis (**Supplementary Fig. S6 A and S6 B**). 1863 identified proteins (1% false discovery rate) were filtered for downstream analysis with a relative standard deviation of less than 25 (**Supplementary Table S3**). The proteins were further analysed for the nature and reproducibility of the transplanted samples wherein we performed a principal component analysis (PCA) of the normalized protein intensity data. The implanted groups showed clear differentiation in PCA space with regards to both the route (SC and O) and the nature of transplantation (syngeneic or allogeneic) due to homology and unique specificity (**Supplementary Fig. S6 C**). The identified proteins also showed excellent reproducibility between replicate runs between the route and nature of implantation (**Supplementary Fig. S6 D**). The N and DC samples of both syngeneic and allogeneic implants also significantly correlated among themselves (**Supplementary Fig. S6 E**) with a dynamic expression pattern (**Supplementary Fig. S6 F**) with ≈88% of proteins common between syngeneic and allogeneic implants suggesting proteome stability (**Supplementary Fig. S6 G**). The expression pattern for both immune response proteins and ECM was unique for the route (SC *vs* O) and nature of the scaffolds (N *vs* DC) implanted.

### Analysis of extracellular matrix protein organization after implantation

The expression of ECM proteins was quite distinct in N and DC implants (**Supplementary Fig. S7 A**). Both N and DC implants shared route-specific homology. ECM proteins were significantly (p<0.05) enriched for GO categories related to proteinaceous extracellular matrix, extracellular matrix organization, disassembly, and extracellular fibril organization (**Supplementary Fig. S7 B**). Furthermore, to determine the functional significance of ECM proteins in syngeneic and allogeneic set-up and to gain new insight into route-specific molecular features, we constructed a functional interaction network with the ECM proteins and visualized in Cytoscape. Four functional modules or clusters were constructed using the MCODE plugin within Cytoscape (**Fig. 3A-D**). It is evident from the cluster organization that ECM components interact with each other to maintain the ECM stability and integrity in the case of both syngeneic and allogeneic implants. In clusters 1, 2, and 4, ECM proteins interact with extracellular space proteins whereas in cluster 3, ECM and extracellular space proteins interact with extracellular fibril organization proteins. Some of these cluster members have a very dense connection and appears to regulate the cluster function. Ranked by degree and other 12 parameters in Cytohubba^25^ and Network Analyzer app within Cytoscape, we identified the top eight proteins (Apoe, Apob, App, Fn1, Kng1, Sparc, Agt, and Actb) as hub proteins along with 4 relevant ECM proteins (Lamb2, Acta2, Ecm1, and Fndc1). Cluster 1 is enriched with five of the eight hub proteins (**Fig. 3A**) whereas cluster 2 and 3 have one each (**Fig. 3B and 3C**). These genes were Apoe, Apob, App, Fn1, Kng1, Sparc, and Agt. The heat map represents the additional hub protein, Actb (**Fig. 3E**). 5 proteins including 3 hub proteins were significantly increased in the N syngeneic SC (App, Sparc, Lamb2 and Fndc1); and allogeneic SC implants (Fndc1). In the O implants, the N allogeneic scaffold showed enhanced expression for 4 proteins (Fn1, Actb, Lamb2, and Ecm1) whereas the N syngeneic implant showed 2 proteins each having either up (Sparc, Agt) or downregulated expression (Fn1, Ecm1). However, the DC syngeneic (LDCO, LDSC) and allogeneic (BNDCSC and BNDCO) implants showed a reduction in the expression of the ECM proteins (**Supplementary Table S3, Supplementary Fig. S7 and S8).**

**Figure 3.**
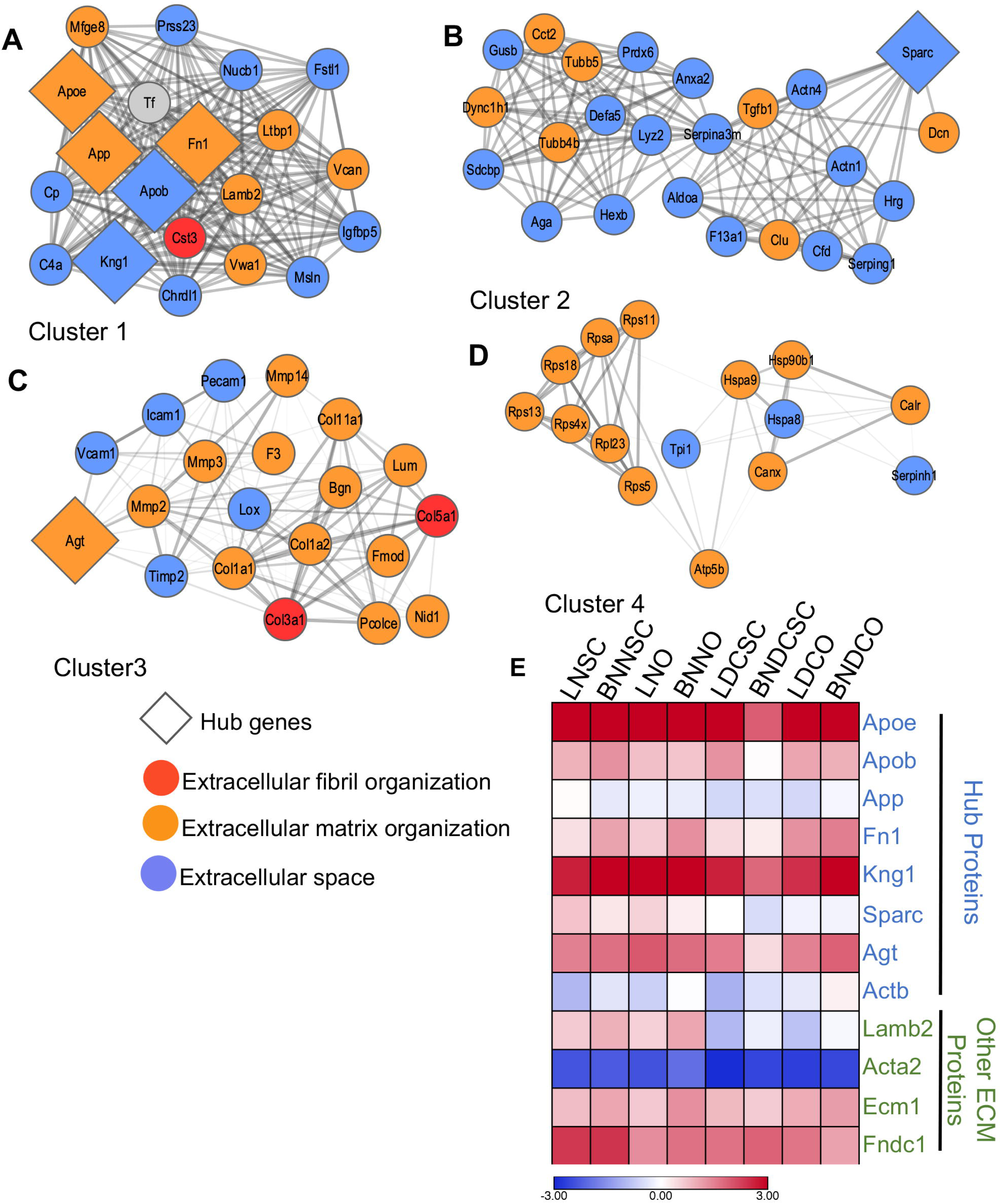
The topological features of the ECM Network of the implants. The PPI network of the 8 hub and related proteins, created by quering the proteins enriched forECM functionagainst the STRING database. The network was exported and visualized usingcytoscape. Proteins were then clustered with MCODE algorithm. Hub proteins were identified using Cytohubba app within the Cytoscape (A-D) MCODE cluster 1-4 respectively. Node colourrepresents gene ontology terms and the edgewidth was proportional to the combined score based on STRING database. (E) Protein expression dynamicsof the hub and selected ECM proteins showing heterogeneity within the transplanted groups. The colour scale represents the log2 normalized expression values.

### Immune protein expression in syngeneic and allogeneic implants

The functional interactions of the immune-related proteins were studied form the proteomic data retrieved from the STRING database. The expression of immune-related proteins was at a lower level in the syngeneic and allogeneic SC implants compared to the O implants. The allogeneic SC implants (N and DC) had significantly less expression level compared to the syngeneic implants (N and DC) for all of the immune-related proteins except Lyn, IL6st, and Cd31 (**Fig. 4)**. On the other hand, the O implants (both N and DC syngeneic and allogeneic) showed more up (C1s, C4, RT1-Aw2, Cd31, Serping 1, Syk and C1qa) than down (Syk, CD4, Lyn, C1qa and C1s) regulation of immune-related proteins **Fig. 4)**. Most of the immune responsive proteins showed dynamic expression involved in the regulation of innate immune responses **(Supplementary Fig. S7 C** and **S7 D**). MCODE analysis revealed the top two clusters for immune-related proteins **(Fig. 4A-B)**.

**Figure 4.**
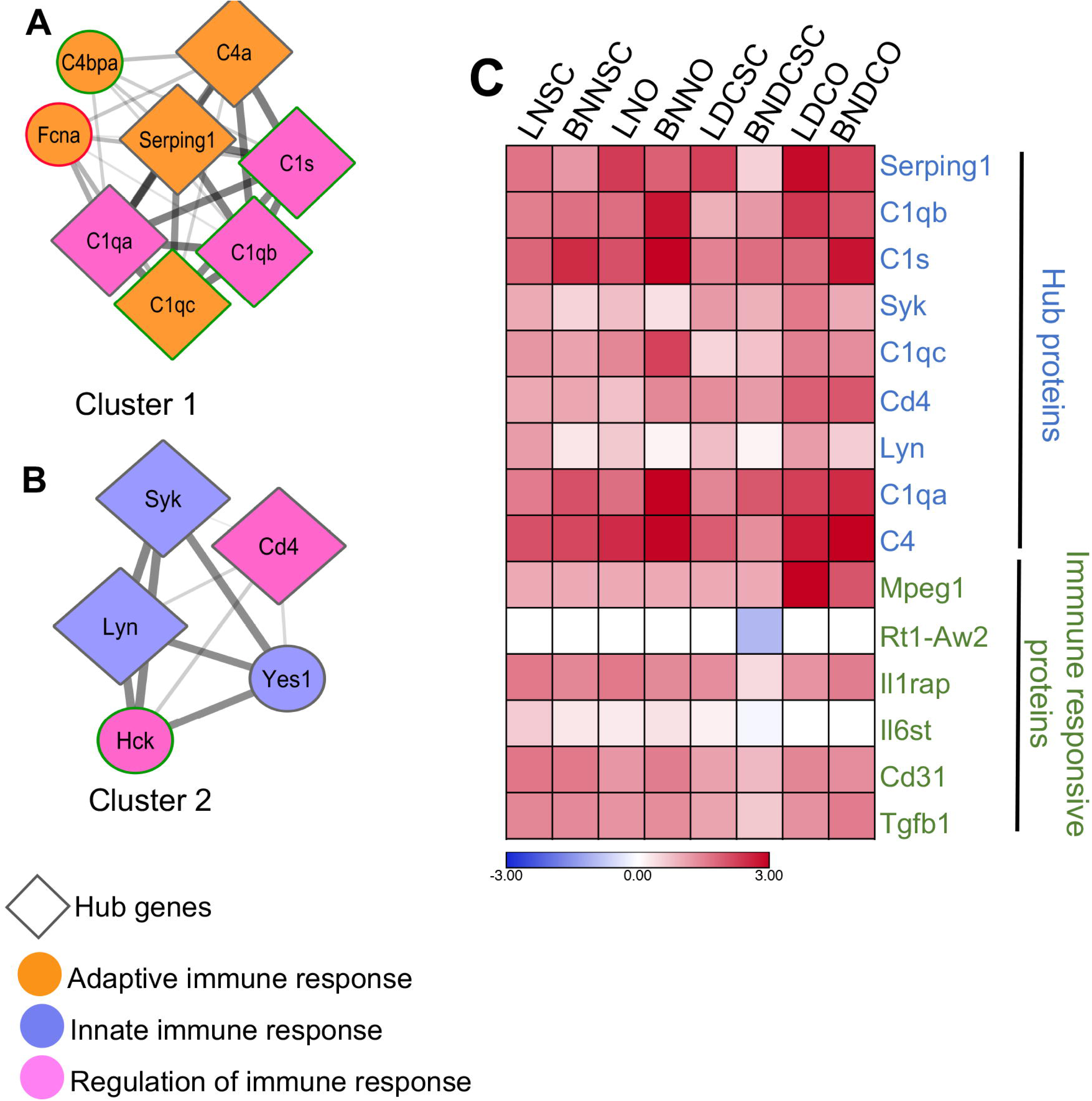
PPI network integration of the immune responsive proteins of the implants. The PPI network was created by quering the proteins enriched for immune function against the STRING database. Network relationships derived from the STRING database were visualized with cytoscape. Proteins were then clustered with MCODE algorithm. Hub proteins were identified using cytohubba app within the cytoscape. **(A, B)** The top 2 MCODE clusters. Eachnode represents a protein and each edge denotes an interaction between a pair of proteins. Node color represents gene ontology terms and the thickness of the edge represents a largercombined-score derived from STRING database. **(C)** Protein expression dynamics of the hub and selected immune responsive proteins showing heterogeneity within thetransplantedgroups. The colour scale represents the log2 normalized expression values.

Cluster 1 is enriched with six of the nine hub proteins **(Fig. 4A)** whereas cluster 2 has three. In cluster 1, proteins with adaptive immune response interact with proteins having the function in the regulation of immune response, which, again interact with innate immune responsive proteins in cluster 2. The 9 significantly upregulated hub proteins, as revealed by Cytohubba and Network Analyser, were Serping1, C1qb, C1s, Syk, C1qc, CD4, Lyn, C1qa, and C4. The other immune markers identified were Mpeg 1, Rt1-Aw2, Il1rap, Il6st, Cd31, and Tgfb1. Out of the 9 significantly expressed HUB proteins, 6 proteins (Serping1, C1qb, C1s, C1qc, C1qa, and C4) were related to the complement system whereas the rest 3 proteins (Syk, CD4, and Lyn) are involved in B and T-cell regulation (**Supplementary Data**). In general, the immune proteins were at an identical level in all implants compared to the normal controls **(Supplementary Fig. S9)**. The number of upregulated and downregulated proteins were identical for the N (6 proteins downregulated and 5 proteins being upregulated) and DC implants (8 proteins downregulated and 5 proteins upregulated) in comparison to the respective N controls (**Supplementary Fig. S7 and S9**).

### Immunohistochemical localization of ECM markers following implantation

Hub proteins from proteomic analysis included Fibronectin (Fn1), Actin (Actb), and Laminin (Lamb2) which have the potential to regulate the cluster function due to their very dense connection which correlates also from bioinformatics predictions. Fibronectin, laminin, and α- SMA which forms an integral component of the ECM were also monitored at the protein level by immunohistochemistry **(Fig. 5**).

**Figure 5.**
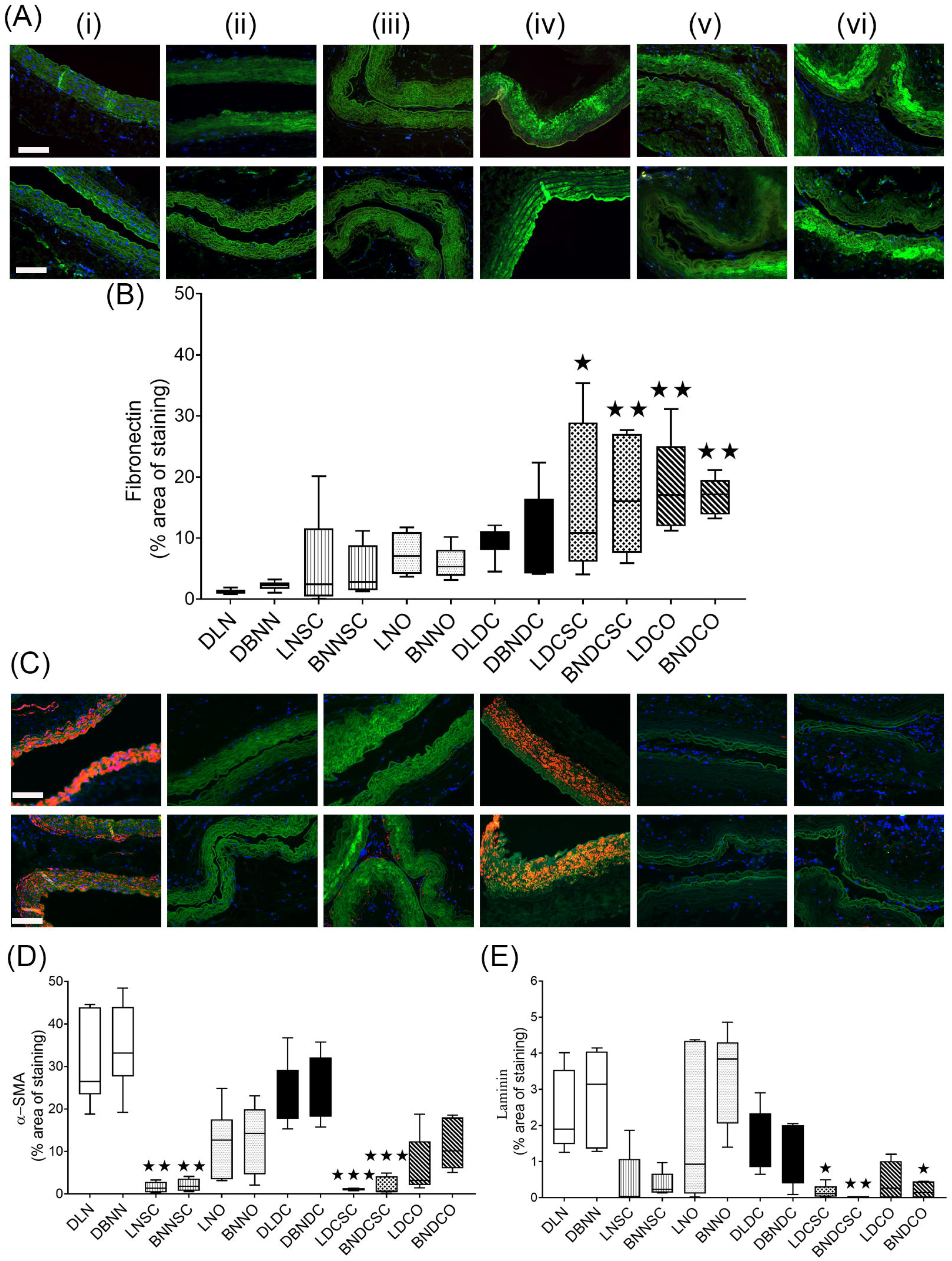
Immunostaining identifying expression of different ECM components in the control scaffolds and implants. Implanted scaffolds after 2 months’ time period were analyzed by histological assessment of different ECMcomponents. The upper panel denotes the implants from Lewis (L) and lower panel from Brown Norway (BN) rats respectively. (i) normal (N), (ii) normal subcutaneous, (iii) normal omentum (NO), (iv) decellularized, (v) decellularized subcutaneous (DCSC) and (vi) decellularized omentum (DCO). A-Localization and B- Quantification of Fibronectin (green colour). C- Localization and D-E- Quantification of α-SMA (red colour) and Laminin (green colour). Representative images are shown at 20x magnification. Scale bars: 100 μm. Results are presented (n=6 in each group) as maximum and minimum of the rank (median shown in the middle of each box). p<0.05 and p<0.01 and p<0.001 indicates a significant difference in comparison to the respective normal implants by Kruskal-Wallis test followed by Dunn’s MultipleComparisons post hoc test.

Post-implantation Fibronectin expression, in the DC SC implants (syngeneic- 13.46 folds, p<0.05; allogeneic- 7.6 folds, p<0.01); DC SC implants (syngeneic- 15.13 folds, ^⍰⍰^p<0.01; allogeneic- 7.68 folds, p<0.01) were significant upregulated compared to the respective N controls (**Fig. 5A and Fig. 5B**). However, all the implants showed reduced expression of α-SMA (**Fig. 5C and Fig. 5D**), compared to the N controls and was significantly lower in both the syngeneic (LNSC-19.65 folds, ^⍰⍰^p<0.01; LDCSC- 27.66 folds, ^⍰⍰^p<0.001) and allogeneic SC implants (N-16.57 folds, ^⍰⍰^p<0.01; and DC- 19.31 folds ^⍰⍰⍰^p<0.001) compared to the N controls. The laminin expression (**Fig. 5C and Fig. 5E**) of the DC implants showed a significant reduction (syngeneic SC-13.68 folds, ^⍰^p<0.05; allogeneic SC-24.15 folds, ^⍰⍰^p<0.01 and allogeneic O-19.65folds, ^⍰^p<0.05) compared to the respective N controls.

### Genomic and protein expression of inflammatory markers following implantation

Since macrophages have been indicated to play an important role in scaffold/implant acceptance, rejection, or modulation, we further characterized the macrophage population at gene and protein level by digital PCR (**Fig. 6)** and immunohistochemistry (**Fig. 7-9**).

**Figure 6.**
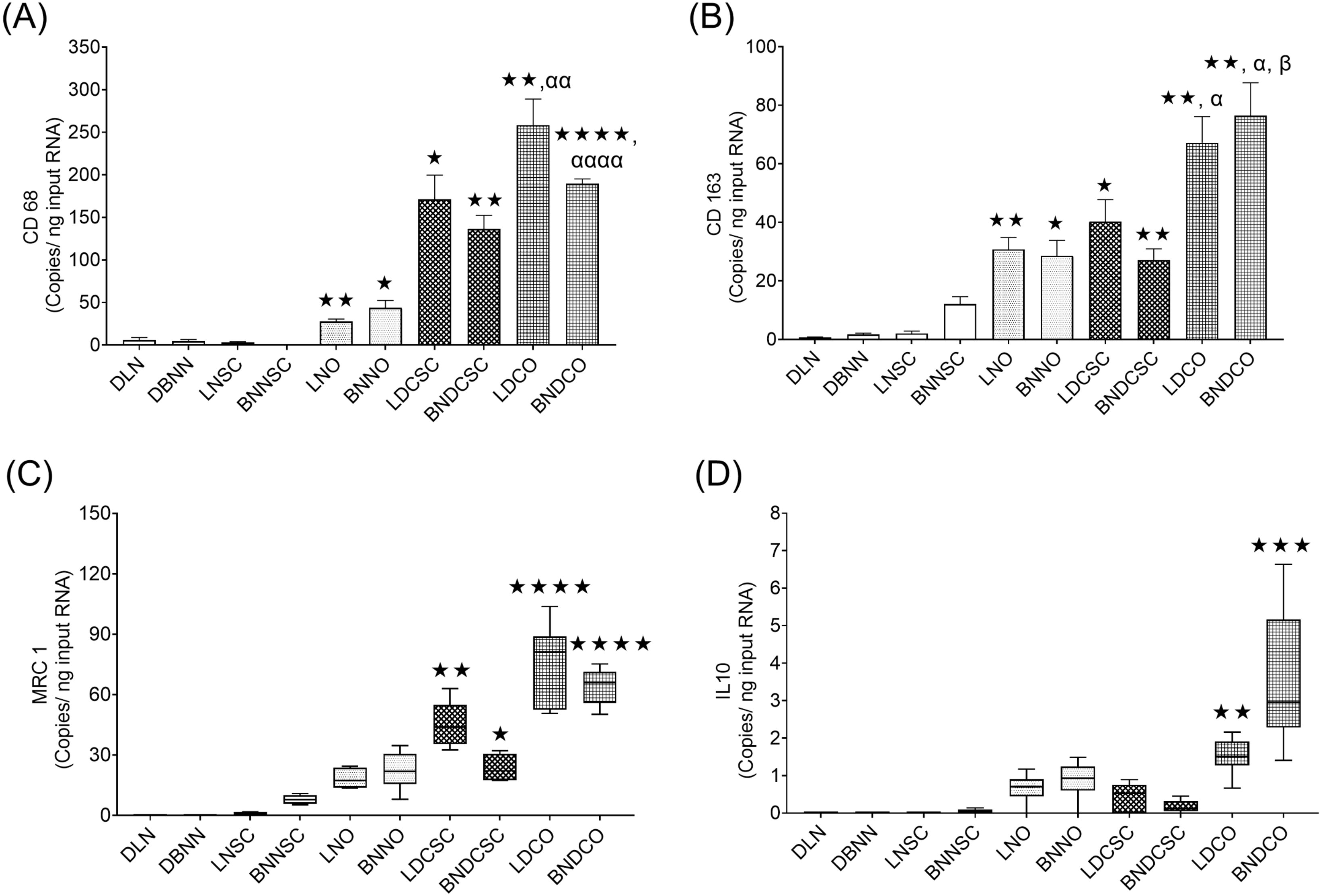
Gene expression analysis of different macrophage markers in the control scaffolds and implants. Implanted scaffolds after 2 months’ time period were characterized for gene expression of different macrophages (M1 *vs* M2 subsets). **A-E**. Copies per ng input RNA of macrophage target gene were calculated using theoretical concentration of the target gene using reference gene, β-actin as outlined in the Materials and methods section. Data are shown as median and range of values (n=6 in each group). **A-B.** Results are shown as Mean±SEM (n=6 in each group). p<0.05, p<0.01, p<0.001 and p<0.0001 indicates a significant difference compared to the respective donor normal scaffold and normal subcutaneous implants; ^α^p<0.05, ^αα^p<0.01 and ^αααα^p<0.0001 indicates a significant difference in compared to the respective normal omentum implants and ^β^p<0.05 indicates a significant difference compared to the respective decellularized implant by One-way ANOVA followed by Dunnett’s T3 MultipleComparisons post hoc test. **C-D.** Results are presented (n=6 in each group) as maximum and minimum of the rank (median in the middle of each box). p<0.05, p<0.01, p<0.001 and p<0.0001 indicates a significant difference compared to the respective donor normal scaffolds by Kruskal-Wallis test followed by Dunn’s Multiple Comparisons post hoc test.

**Figure 7.**
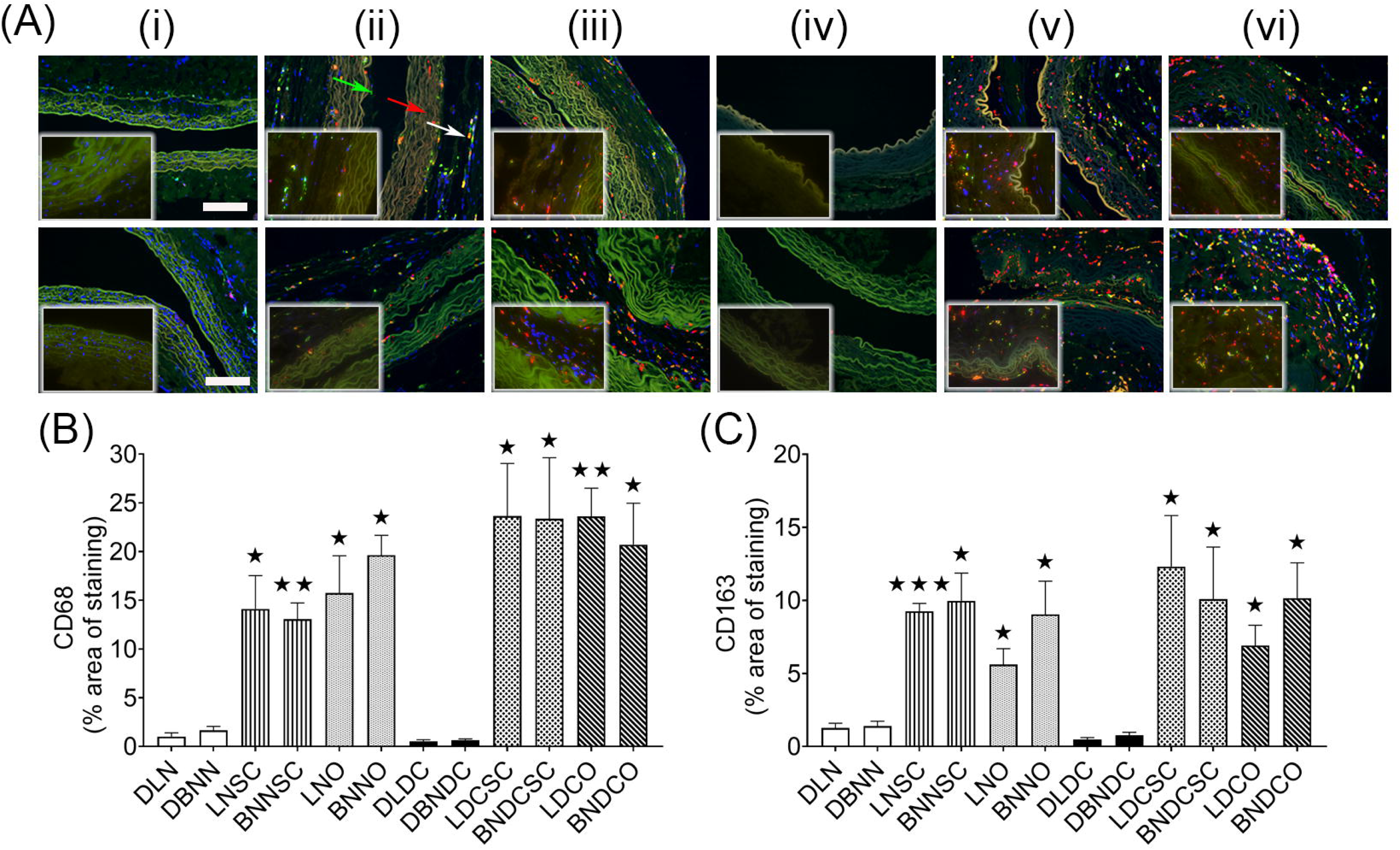
Immunostaining identifying macrophage markers, CD68 and CD163 in the control scaffolds and implants. Implants after 2 months’ time period were subjected to measurement of macrophage (CD68 and CD163).The upper panel denotes the implants from Lewis and the lower panel from Brown Norway rats respectively.**Arrows**- red:CD68 and green: CD163. **7A**- Localization and **7B-7D**- Quantification of CD68 and CD163 macrophages. Data are presented as Mean±SEM (n=6 in each group). Representative images are shown at 20x magnification. Scale bars- 100 μm. p<0.05, p<0.01 and p<0.001 indicates a significant difference in comparison to the respective donor normal scaffolds by One-way ANOVA followed by Dunnett’s T3 Multiple Comparisons post hoc test.

**Figure 8.**
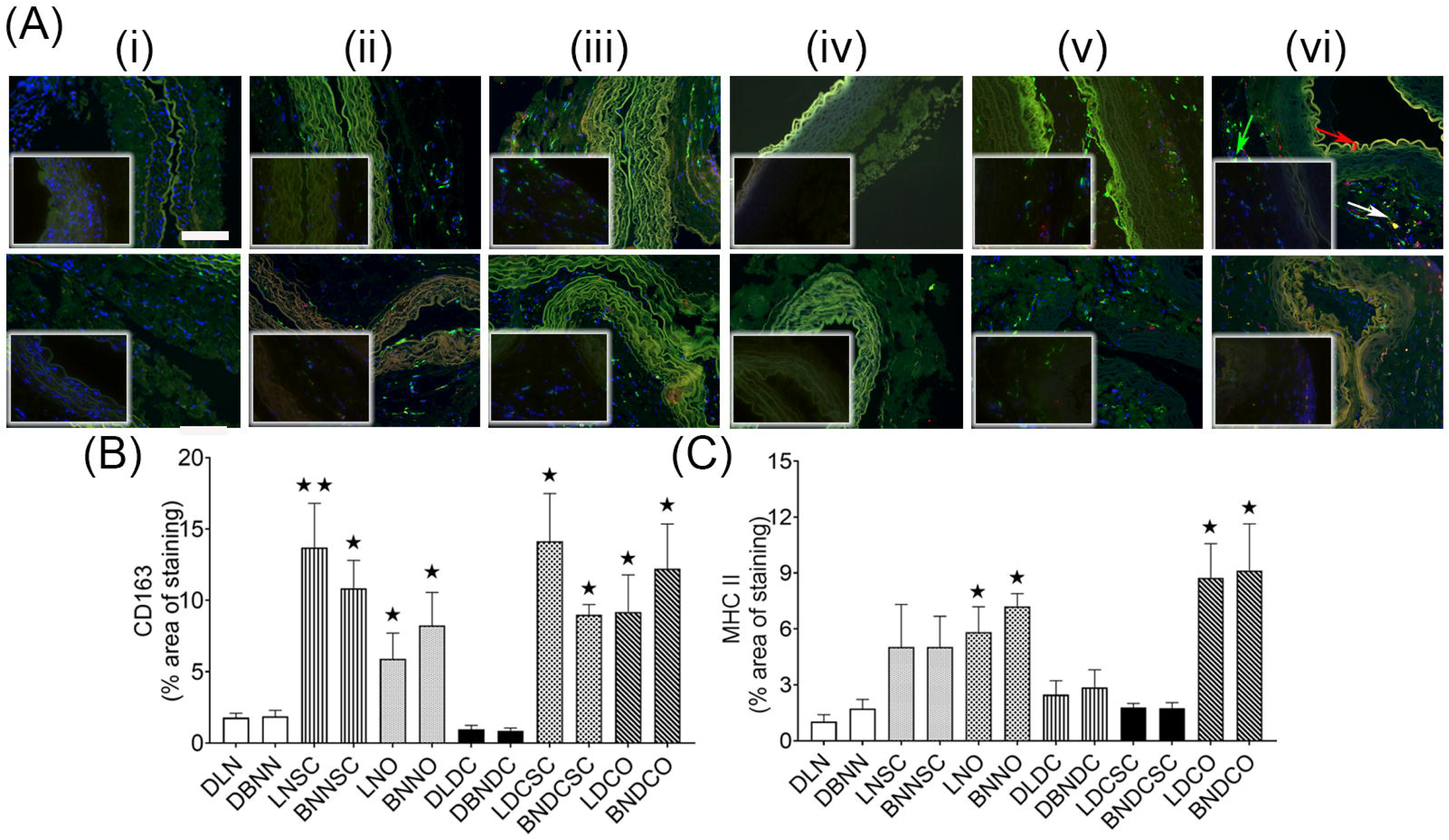
Immunostaining identifying macrophage markers, CD163 and MHCII in the control scaffolds and implants. Implanted scaffolds were subjected to immunofluorescence assessment for different macrophage (CD163 and MHC II) as outlined in Materials and Methods section. The upper panel denotes the implants from Lewis and the lower panel from Brown Norway rats respectively. **Arrows**- red: MHC II and green: CD163 **8A**- Localization and **8B-8D**- Quantification of CD163 and MHCII macrophages. Representative images are shown at 20x magnification. Scale bars: 100 μm. Data are presented as Mean±SEM (n=6 in each group). p<0.05 and p<0.01 indicates a significant difference in comparison to the respective normal implants by One-way ANOVA followed by Dunnett’s T3 Multiple Comparisons post hoc test.

**Figure 9.**
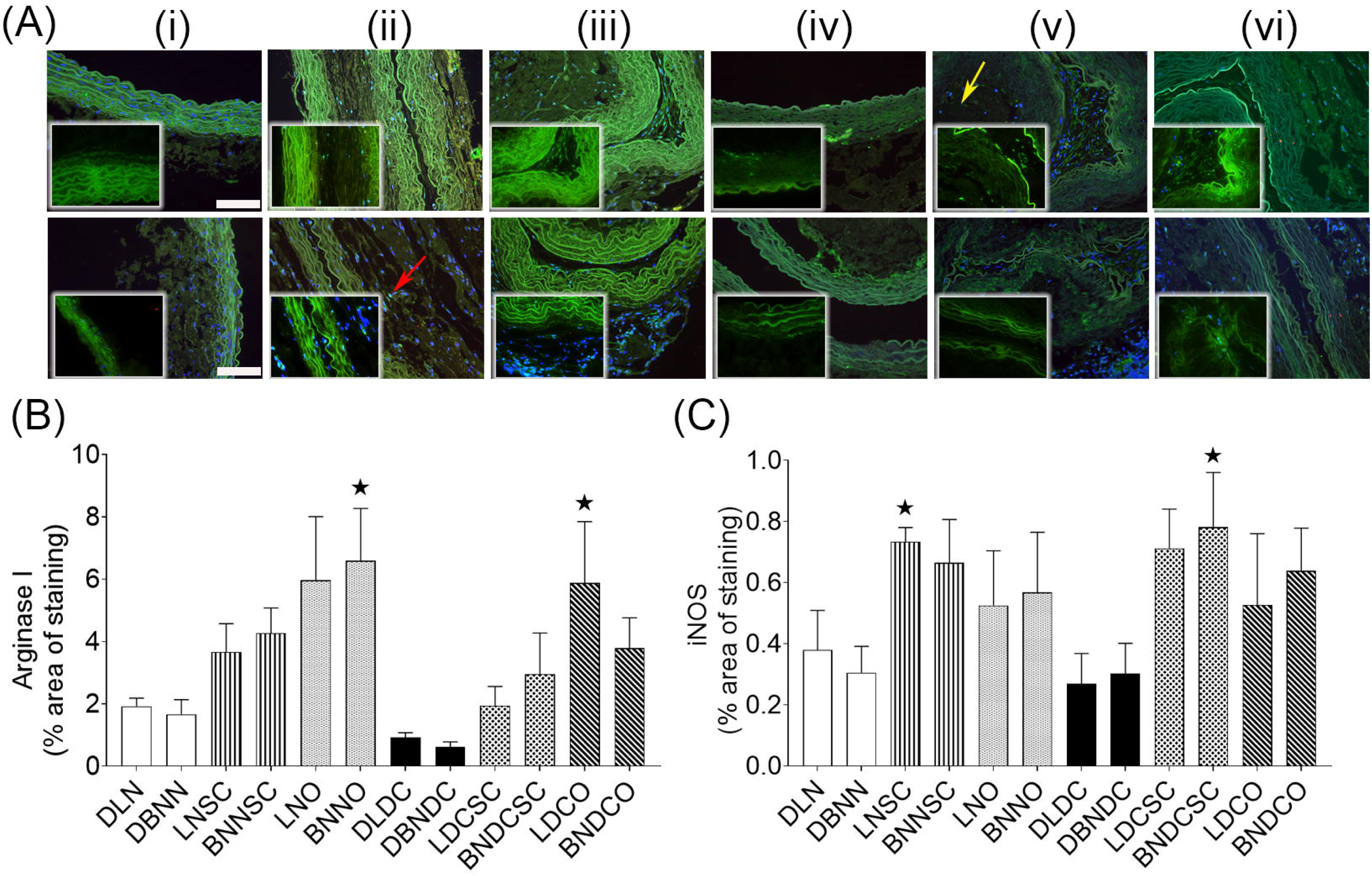
Immunostaining identifying macrophage markers, Arginase I and iNOS participating inthe control scaffolds and implants. Implanted scaffolds were subjected to immunohistological assessment for different macrophage (Arginase I andiNOS). The upper panel denotes the implants from Lewis and the lower panel from Brown Norway rats respectively. 9A- Localization and 9B-9C- Quantification of Arginase I and iNOS macrophages. **Arrows**-red: iNOS and green: Arginase I. Representative images are shown at 20x magnification. Scale bars: 100 μm. Data are presented as Mean±SEM (n=6 in each group). p<0.05 indicates a significant difference in comparison to the respectivenormal scaffolds by One-way ANOVA followed by Dunnett’s T3 Multiple Comparisons post hoc test.

The genomic expression pattern of the macrophage markers- CD68, CD163 and MRC1 showed significant expression in all the implants (**Fig. 6A-D**). The internal reference was β actin gene and the target gene expression were done as outlined in **Supplementary file** and **Supplementary Fig. S10.**

The gene expression level in the N controls was at a very low level and were expressed in an order CD68>CD163>MRC1≈IL10. Also, the CD68 and CD163 gene expressions of the N SC donor implants were identical to that of the N controls irrespective of the genetic backgrounds. For the CD68 expression, the N O implants (syngeneic- 4.74 folds, p<0.05; allogeneic-10.09 folds, p<0.01) showed significantly higher expression compared to the N controls and N SC implants (**Fig. 6A**). All the DC implants both syngeneic (SC, - 28.980 folds, p<0.05 and O- 43.76 folds; p <0.01) and allogeneic (SC- 31.47 folds, p <0.01and O- 43.73 folds; p <0.0001) implants showed significantly higher expression compared to their respective N controls and N SC implants **(Fig. 6A).** Furthermore, the DC O implants (syngeneic - p <0.01 and allogeneic - p^⍰⍰⍰⍰^<0.0001) had significantly higher expression compared to the N O implants.

The CD163 gene expression in case of the N O implants (syngeneic - 43.92 folds, p <0.01 and allogeneic - 17.11 folds; p <0.05) was significantly higher than the respective N controls. The DC SC implants (syngeneic - 57.49 folds, p<0.05; allogeneic- 16.24 folds; p <0.01); DC O implants (syngeneic - 95.92 folds and allogeneic- 45.72 folds; p<0.01) showed significantly higher expression compared to the N controls and SC implants. Also, the DC O implants showed higher expression ( p<0.05) compared to the N O implants and DC SC implant (allogeneic, p<0.05) **(Fig. 6B)**.

The level of MRC-1 gene expression was very negligible in the N controls and was not significantly increased in case of N implants. However, all the DC implants showed higher expression **(Fig. 6C)**. The MRC1 expression was at a significantly higher level in the DC SC (syngeneic, ^⍰⍰^p<0.01 and allogeneic, ^⍰^p<0.05) and DC O (syngeneic, ^⍰⍰⍰⍰^p<0.0001 and allogeneic, p<0.0001) implants compared to the N controls **(Fig. 6C).**

The anti-inflammatory cytokine, IL 10 expression was negligible in case of N controls and N SC implants. The expression progressively increased in the order N O implant < DC SC implants < DC O implants. The DC O implants showed significant increase (syngeneic- p^⍰⍰^<0.01 and allogeneic- p^⍰⍰^<0.001) compared to the respective N controls (**Fig. 6D**).

Targeted multiplexing of two protein targets by immunohistochemistry was used to detect (i) CD68 (pan macrophage) and CD163 (M2 macrophage); (ii) MHCII (M1 macrophage) and CD163 (M2 macrophage) and (iii) iNOS (M1 macrophage) and Arginase I (M2 macrophage) in the N and DC controls, and SC and O implants by immunohistochemistry (**Fig. 7-9) and Supplementary Fig. S11)** using validated antibodies and appropriate positive control staining **(Supplementary Fig. S12).**

The expression of the pan, M1, and M2 macrophage markers was determined by dual staining as described under the **Supplementary file**. The percentage of positive cells for a particular marker was expressed in terms of the nucleated cells covering particular staining area. For the pan macrophage marker, CD68 the DC implants (≈14-23 folds; p^⍰^<0.05- p^⍰^<0.01) and the N implants (≈8-15 folds; p^⍰^<0.05) showed significant expression compared to the N controls (**Fig. 7A and 7B**).

The CD163 expression paralleled the CD68 expression albeit at a lesser level (≈2-3 folds lesser) compared to the N controls. However, there was almost identical CD163 expression of different implants (N *vs* DC) placed at different sites (SC *vs* O) (**Fig. 7A and 7C**).

Subsequently, we also monitored the expression of major histocompatibility antigen, MHCII as a potential M2 marker in the different implants along with CD163 expression (**Fig. 8**).We could detect significant increase of MHC II macrophages in the N O implants (4.17-5.67 folds, p <0.05) DC O (8.511 folds and 5.295 folds, p <0.05) implants compared to the N controls (**Fig. 8A and 8C**).

We also performed immunolabelling for two additional macrophage markers, iNOS (M1 macrophage) and Arginase I (M2 macrophage) (**Fig. 9**). Significant increase in expression of M2 macrophage, Arginase I (**Fig. 9A-9B**) was found mainly in the N allogeneic (3.96 folds, p <0.05) and DC syngeneic (3.06 folds, p <0.05) implants. The expression of the M1 macrophage, iNOS was however weak **(Fig. 9A-9C)** and was mainly observed in the SC (syngeneic N SC- 1.93 folds, p <0.05 and allogeneic DC - 2.56 folds, p <0.05) implants compared to the N controls.

Histological observations of allogeneic and syngeneic implants revealed mainly lymphocyte infiltrations **(Fig. 2 and Supplementary Fig. S2)**. Subsequently, we also monitored T-cells (CD3^+^, CD4^+^, and CD8^+^) infiltration in the implants by immunohistochemistry which was mainly found around the implants with only few cells present in the lumen of the implants (**Fig. 10**).

**Figure 10.**
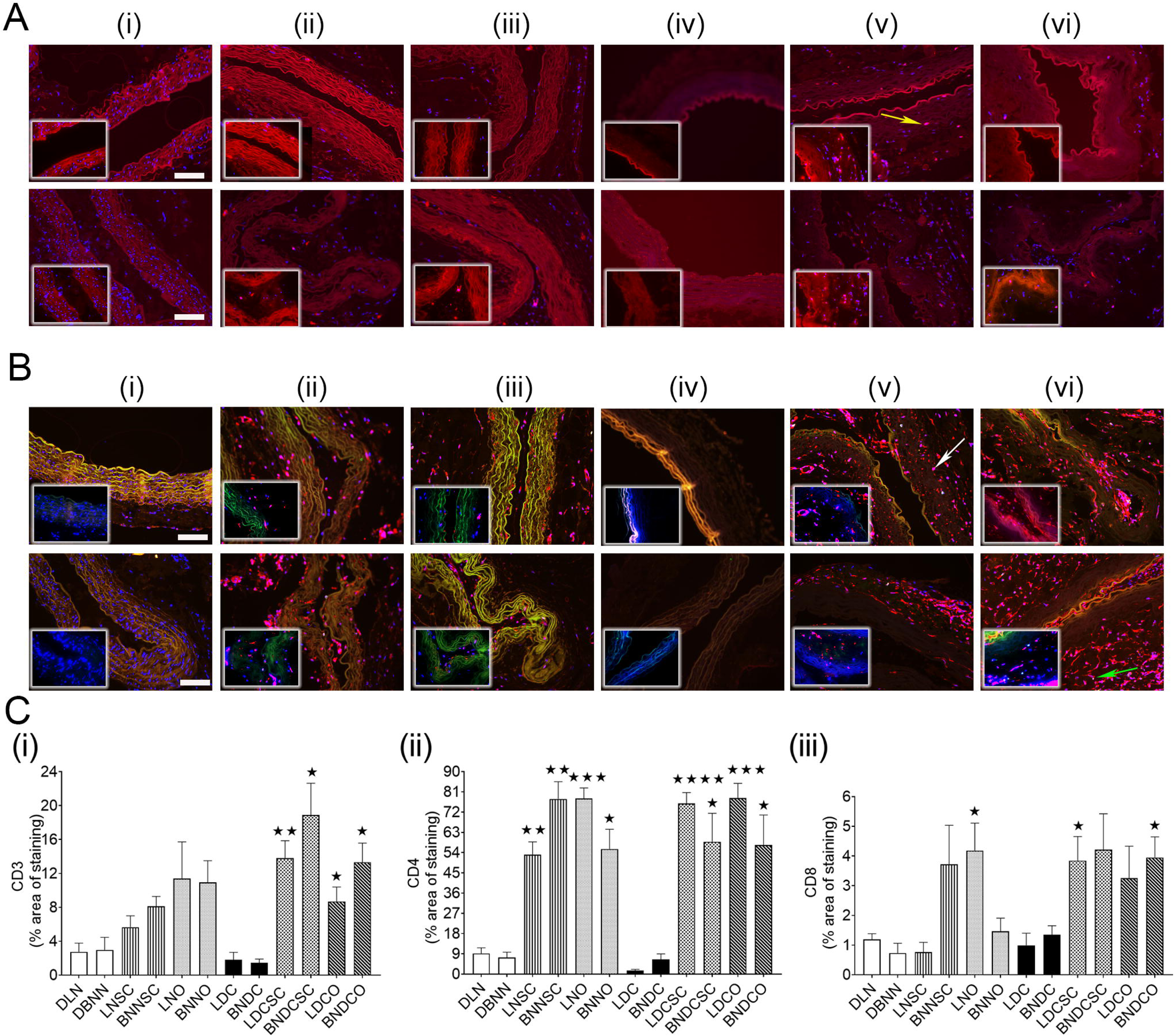
Immunostaining for immune markers, CD3, CD4 and CD8 in the control scaffolds and implants. Implanted scaffolds after 2 months’ time period were harvested from the subcutaneous and omentum site and analyzed by immunofluorescence assessment for different immune markers (CD3+, CD4+ and CD8+cells). The upper panel denotes the implants from Lewis and the lower panelfrom Brown Norway rats respectively. Yellow arrow denotes the presence of CD3+ve cells, white and green arrow denotes the presence of CD4+ve and CD8+ve cells respectively. Localization **(10A)** and quantification of CD3 (red colour) (**10C i)**; Localization **(10B)** and quantification of CD4 (red colour) and CD8 (green colour) (**10C ii-iii)** is shown. Representative images are shown at 20x magnification. Scale bars: 100 μm. Data are presented as Mean±SEM (n=6 in each group). p<0.05, p<0.01, p<0.001 and p<0.001 indicates a significant difference in comparison to the respective normal implants by One-way ANOVA followed by Dunnett’s T3 Multiple Comparisons post hoc test.

The expression of the pan T-cell marker, CD3+ was not significantly increased in the normal syngeneic and allogeneic implants. All the DC implants from the SC and O route showed significant upregulation of the CD3+ expression compared to the respective N controls (syngeneic DC SC- 5.03 folds, ^⍰⍰^p<0.01, syngeneic DC O- 3.17 folds, ^⍰^p<0.05; allogeneic DC SC -6.36 folds, ^⍰^p<0.05 and allogeneic DC O -4.48 folds, ^⍰^p<0.05) (**Fig. 10A and 10C i**). Of the T-cell markers, CD4+ and CD8+, CD4+ was the more dominant T-cell populations (**Fig. 10B and 10C ii-iii**). There was varied expression of syngeneic (N SC- 5.77 folds, ^⍰⍰^p<0.01; N O-8.48 folds, ^⍰⍰⍰^p<0.001; DC SC- 8.24 folds ^⍰⍰⍰⍰^p<0.0001; DC O-8.49 folds, p<0.001) and allogeneic (BNNSC-10.5 folds, p<0.05; BNNO- 7.50 folds, p<0.05; BNDCSC- 7.94 folds, ^⍰^p<0.05; and BNDCO- 7.74 folds, ^⍰^p<0.05) implants compared to the N controls (**Fig. 10B and 10C ii**). CD8+ expression was at a much lesser level only in few groups (syngeneic N O, syngeneic DC SC and allogeneic DC O-3.2-.4 folds, ^⍰^p<0.05) compared to the N controls (**Fig. 10B and 10C iii**).

The systemic immune response following implantation was identical in all the recipient rats and were also similar to that of the sham treatment at the end of 2 months (**Fig. 11A and 11B)**. The expression pattern of cytokines is probably indicative of the absence of antibody-mediated rejection following scaffold implantation.

**Figure 11.**
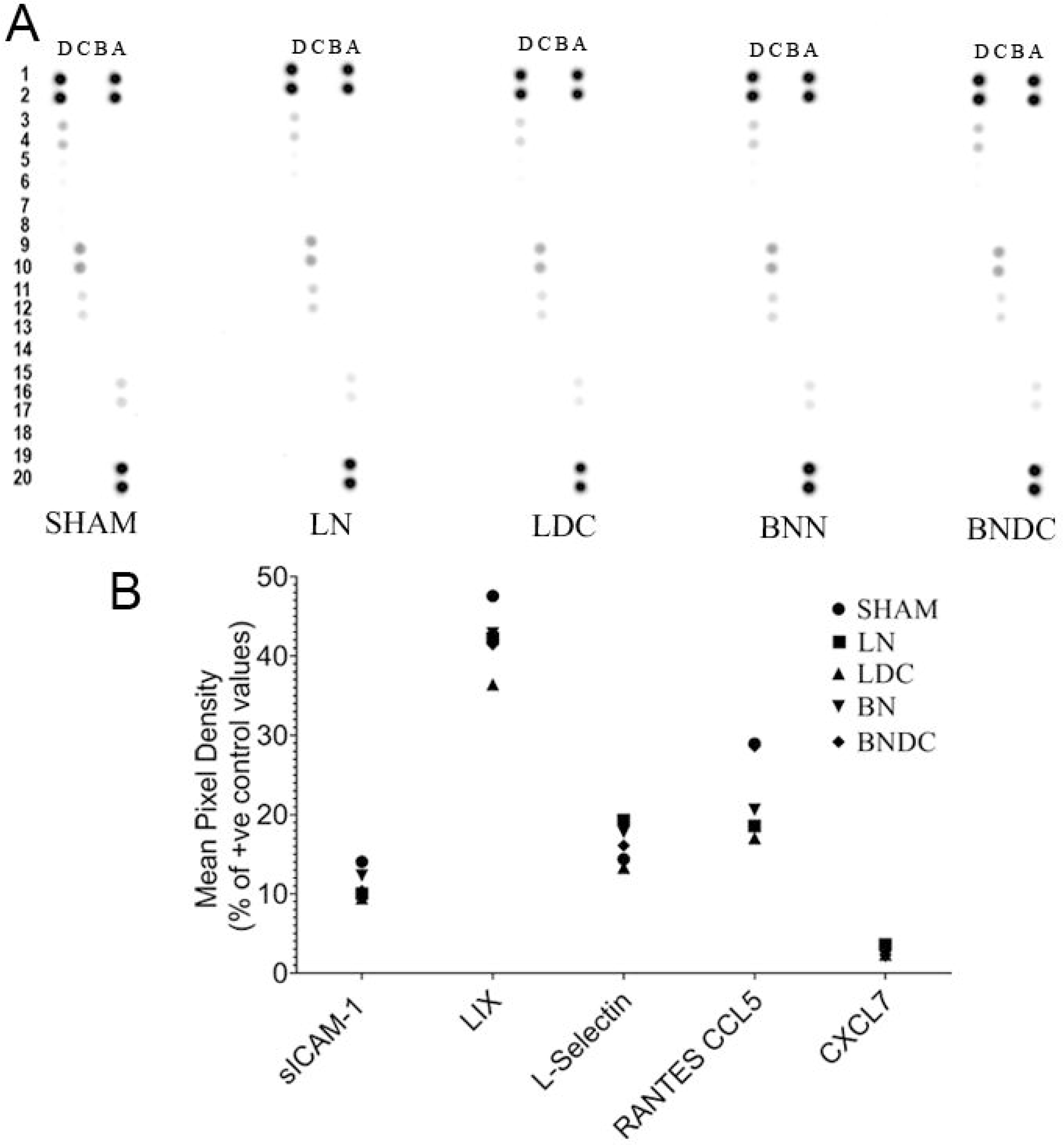
Cytokine and chemokine detection in serum of recipients and SHAM group. A. Membrane based proteome profiling of selected cytokines and chemokines. Serum from implanted recipient Lewis female rats (LN, LDC, BNN and BNDC) was compared with the SHAM operated Lewis female rats and analysed for the relative level of analytes as specified in the Materials and Methods section. B. The calculated level of selected analytes showing most prominent expression for all the groups. The relative level of cytokines and chemokines was expressed as Mean Pixel Density after suitable correction of negative and positive reference spots present in the array membrane.

## Discussion

The current study compared blood vascular scaffold prepared by chemical decellularization with that of acellular scaffold prepared *in vivo* at genomic and protein level. The development of an acellular scaffold by natural means could help overcome the problem associated with synthetic, biological scaffolds as well as allogeneic scaffolds possibly minimizing of unfavourable immune responses due to selection of appropriate routes.

During the study, we initially prepared scaffold initially by chemical decellularization using an alternate cycle of SDC and Glycerol/EDTA mix^20^ which showed loss of tissue architecture indicative of the deleterious effect of chemical decellularization.^26, 27^ To monitor the fate of syngeneic and allogeneic implants following *in vivo* implantation, normal untreated (N) and chemically DC/acellular scaffolds were placed as a free-floating implant without vascular anastomosis at subcutaneous (SC) and in omentum (O). The implants (SC and O) behaved uniquely according to the route of implantation as evidenced from the genomics and protein characterization. The O implants irrespective of the genetic background were found in general to have a poor outcome compared to the SC implants as assessed from the architecture, fibrosis and necrosis score. The outcome was further worsened if the scaffold was chemically DC prior to implantation. The presence of lymphocytes in the implants suggests the ensuing immune response to be of chronic type. However, minimum inflammation associated with the implants particularly the N implants (except the allogeneic O implants) is suggestive of a favourable modulation of these implants *in vivo*.

ECM and immune responsive protein dynamics following implantation were monitored by quantitative proteomics and immunohistochemistry. The proteomics findings were evaluated further by protein interaction network which indicated the involvement of unique ECM proteinclusters operating at extracellular space, involving extracellular fibril organization proteins. The difference in nature of the scaffold and route of implantation was reflected from the hub ECM and immune proteins expression with the subcutaneous implants showing more uniform expression of ECM and immune-related proteins compared to the O implants. The Extracellular Matrix Protein 1 (Ecm1)was up-regulated mainly in N tissues irrespective of origin (syngeneic *vs* allogeneic). Similarly, the Fibronectin type III domain-containing protein (Fndca), which is associated with G protein signaling was also upregulated in general. However, the ECM proteins like Fibronectin 1 (Fn1) and laminin (Lamb2) were found to be upregulated in N controls whereas the expression was reduced in DC controls. Subsequently, immunohistochemical tool further showed the expression pattern of three important ECM proteins, Fibronectin, α-SMA and laminin. Except for fibronection, the other two ECM proteins, α-SMA and Laminin did not show increase in the expression following implantation. However, the DC implants showed enhanced expression of fibronectin compared to other implants. A marked increase in the expression of ECM components e.g. vascular laminins and fibronectin is a common phenomenon during early syn- and allogeneic transplantation and may trigger lymphocyte recruitment at the implantation site.^28^ This may be the reason for the association of immune response with most of the implants studied. Also, Alpha actin (Acta2) or α-SMA which is normally associated with chronic rejection in all scaffolds^29^ was not up-regulated in any of the implants. The deposition of a fibrous capsule around DC ECM based scaffolds which normally elicits immune response limiting tissue remodelling and regenerative potential^6^, was mainly limited to the normal omental implants with additional necrosis also observed for the DC O implants due to a probable unresolved inflammatory response. However, the chemical DC process used by us mostly preserved the ECM components as seen from the immunohistochemical data. Following DC, however, the ECM epitopes got exposed to the antibodies due to the lack of cells and were probably the reason for the higher expression level compared to the normal implants. Any additional increase in the ECM protein expression could be attributed to the post-implantation modifications. The proteomics findings pointed to the expression of the immune proteins particularly the innate arm of the immune response with overlapping roles with the adaptive immune system. The immune markers detected were normally associated with complements, macrophages, and immune cells category with hub immune proteins upregulated in almost all the implants. However, some of the hub proteins were expressed preferentially in omentum compared to the subcutaneous implants e.g. Serping1, C1qb, C1s, C1qc, C1qa, and C4, suggestive of a possible counterbalance immune response in an omental milieu compared to the subcutaneous site. Anti-HLA class I antibodies are primarily implicated in acute rejection, whereas anti-HLA class II antibodies during late rejection. The protein expression of MHC I molecules (RT1-Aw2) and also MHC II in syngeneic and allogeneic implants were identical. The complement proteins can recognize danger signals generated by foreign particles e.g. intruding pathogens or danger-associated molecular patterns (DAMPs) presented by transplanted scaffolds.^30^ Complement component, e. g. Syk, CD4, and Lyn plays important roles in adaptive, and innate immune response as part of the host defence system. CD4, which is a membrane glycoprotein and a member of the immunoglobulin supergene family acts as a co-receptor in MHC class II-restricted T-cell activation by direct and indirect allorecognition.^31^ The tissue macrophages are one of the effector cells against injury and scaffold implantation led to modulation of their own phenotype and surrounding cells. M1 macrophages are characterized by the high expression of MHC II and inducible nitric oxide synthase (iNOS) apart from the pan macrophage marker, CD68 whereas M1 macrophages are characterized by M2-associated genes such as arginase-1 (Arg1), Mrc1, CD163, and (TGF-β1).^32^ We focused on the identification of mainly two broad phenotypes of macrophages - M1 (pro-inflammatory) and M2 (anti-inflammatory) which are reported to regulate a host of inflammatory cytokines following implantation. Macrophages, which are an integral part of innate immune system, based on stimulation are classified as M1 (stimulated with interferon gamma, IFN-γ/lipopolysaccharide, LPS) and M2 (stimulated with interleukin-4, IL-4/interleukin-13, IL-13) macrophages. With genomic analysis we detected pan, M1 and M2 macrophage subsets (CD68- pan macrophage, CD163-M2 macrophage, MRC-1-M2 macrophage, and iNOS-M1 macrophage). Proteomics study revealed normal omentum (NO) and DC implants to have significant protein expression which could possibly explain the up regulation of protein expression seen in subsequent immunohistochemical analysis. Interestingly, two M2 macrophage lineage genes (CD163 and MRC1) and the anitinflammatory cytokine, IL-10 expression was found to be at the highest level in the implants where the modulation, i.e. syngeneic and allogeneic DC O implants. We could also identify infiltrating macrophages and their phenotypes (CD68 -pan), MHCII -M1, iNOS -M1, CD163 -M2 and Arginase -M2 as a putative immune response after implantation. The macrophage infiltration was predominantly located at the muscular area with fewer cells towards the lumen. The N and DC O implants showed higher macrophage infiltration compared to the N control and SC implants. The pan macrophage marker, CD68 as well as the M2 macrophage, CD163 expression was uniform across all the syngeneic and allogeneic implants at all routes. The expression of the M1 macrophage, MHCII was more prominent with the O implants whereas for M1 macrophage, iNOS there was no uniform expression. The higher expression of MHC II in O implants is possibly not a favourable sign of graft acceptance which is also mirrored earlier in the histopathological changes. The level of expression for the M2 macrophage, Arginase I was also more in the O implants. The critical role of M2 macrophages in the remodelling of engineered vessels after implantation due to the promotion of blood vessels, cell survival, and tissue regeneration has earlier been demonstrated.^33, 34^ Even though M1 macrophages has been associated mostly with the O implants, the expression of macrophages particularly the M2 subsets, CD163, Arginase I and MRC 1 could possibly indicate the type of immune response at the implantation site, a possible effort in *in vivo* to facilitate favourable remodelling of the implanted scaffolds without rejection.

An elevation of inflammatory markers and immune cell circulation has been reported to be one of the main causes of all scaffold rejection promoted us to characterize tissue distribution of T-lymphocytes after implantation. To monitor the expression of the T-cell mediated rejection of implanted tissue, the presence of CD4+, CD8+ lymphocytes were detected along with the pan T-cell marker, CD3+ cells. Earlier CD4 protein was also detected as a hub protein by proteomics. Moreover, the CD4+/CD8+ ratio is usually increased after solid allogeneic/alloscaffold transplantation and is an important index for evaluating immune rejection.^35, 36^ Our study showed that the CD4+ and CD8+ cells in the syngeneic and allogeneic scaffolds were almost at an identical expression level. The presence of identical immune marker having the identical expression level after implantation of syngeneic and allogeneic scaffolds suggests the involvement of a sterile inflammation at the implantation site which is produced irrespective of the genetic backgrounds. A statistically significant increase in the CD3+ cell was observed in all the DC implants whereas an increase of CD4+ cells with a minimum infiltration of CD8+ cells was observed in both the N and DC implants, suggestive of an absence of a strong T-cell-mediated immune response. The cellular infiltration reveals the nature of the T-cell infiltrate which is mainly CD4+, suggestive of the absence of a rejection process which opens up the possibility to create an acellular biological scaffold with all essential ECM components maintained. Both the syngeneic and allogeneic implants showed well preserved smooth muscles of blood vascular architecture and minimum intralumen cellular infiltrate with predominant mononuclear cell infiltration confined to the periscaffold space. Statistically significant increased proportion of CD4+ T-lymphocytes in the implanted scaffolds observed could lead to improved scaffold outcome after implantation as has been associated with allogeneic transplant acceptance.^37^ The immune responsive proteins were mostly upregulated in N allogeneic O implants suggestive of a more localized immune response in the omentum. In syngeneic transplants, the pattern was quite distinct, with N syngeneic transplants showing upregulated immune responsive proteins in comparison to the DC syngeneic implants with down-regulated immune response proteins. We have seen earlier from gene expression study the increased expression of the anti-inflammatory cytokine, such as IL-10 in syngeneic and allogeneic DC O scaffolds where there was consistent macrophage infiltration. The presence of macrophage infiltration is suggestive of a possible protection from tissue damage caused by excessive inflammation because of potential crosstalk with a subpopulation of T cells such as regulatory (Tregs).^38, 39^ The presence of CD31, (PECAM 1) normally expressed on early and mature vascular endothelial cells, observed during our proteomic analysis could be an indication of repopulation with recipient non-immune cells. However, endothelial cells also act as site for leukocyte adhesion rendering them a potent immunostimulatory inflammatory components in response to transplanted tissues. Loss of endothelium as seen during histology-based assessment possibly could lead to higher incidence of thrombus formation on one hand but reduce the antigen presence and adhesion molecule expression on the other thus possibly minimizing inflammatory response against the graft^40^ as seen during our study. The endothelial lining was removed to a large extent during the histological processing and was absent in the decellularized scaffolds and the implants.

In terms of the systemic immune response, there were no serum markers suggestive of any systemic inflammatory responses. All the detected immune markers were found to be at the same level as the sham treated rats supporting a more favourable outcome of scaffold remodelling. We observed immune responses which did not alter the scaffold architecture after implantation. This observation probably could be attributed to the anatomical feature of the aorta having a large diameter, the rigidity of the vessel as well as the implantation route selected. The SC site allowed minimizing the ensuing immune responses whereas the omentum prompted a more vigorous immune response which leads to alteration in the tissue architecture of the scaffold. The chemical DC strategy used was aimed to minimize the tissue architecture damage in terms of the ECM or triggering of an adverse immune reaction, which was to some extent again circumvented by choosing the SC route for implantation. The host response at different sites was similar to non-resorbable foreign material characterized by low-grade chronic inflammation, negligible scaffold degradation and or necrosis and fibrosis was supported by our histology and immunohistological findings. The differential responses of tissue scaffolds placed SC^41^ and in the omentum^42^ have been reported earlier and points to the presence of innate immune cells particularly macrophages in the scaffold which could have contributed to the acellular state of both the syngeneic and allogeneic scaffolds in our study.

Biological scaffolds based on ECM have been shown to normally promote a macrophage switch from M1 to M2 cell population deciding the outcome of post-implantation. However, chemical DC can expose the ECM preventing the beneficial M2 response which can lead to scar tissue formation.^43, 44^ Moreover, alteration of the tissue-specific ECM composition following organ transplantation has also been reported.^45^

So, we decided to compare an acellular scaffold/implant prepared *in vivo* without the involvement of chemicals but rather harnessing the body’s natural immune defense system, i.e. predominantly the innate arm of the immune system i.e. macrophages and partly adapted immunity through the involvement of CD4^+^ T-helper cells. From our histological findings, we could conclude that there was mostly maintenance of the typical smooth muscle architecture as well as the absence of a fibrous capsule around the N and DC implants. However, we did not evaluate the N or DC implants for further vascular anastomosis but rather used them as a free-floating scaffold to evaluate the alteration of the scaffold/ECM and ensuing immune responses after implantation. A functional evaluation along with the dynamic observations of the implanted scaffolds is a future option that we would like to explore.

## Conclusion

The fate of biological grafts prepared by chemical and natural decellularization utilizing different genetic backgrounds of donor and recipient mice after implantation have been studied at protein and molecular level in the current study. The study was carried out without using any immunosuppressant allowing dynamic observation of the implants under a normal immunological set up. The study opens up the possibility to utilize body’s immune machinery to produce an acellular scaffold that has the potential to support cellular growth due to the maintenance of ECM components as well as triggering of identical immune reactions in both syngeneic and allogeneic implants. The importance of the development of a natural biological scaffold also stems from the concern of the fast commercialization of tissue-engineered organs e.g. decellularized tissue for clinical application in humans by FDA and CE which could possibly jeopardize patient’s safety on the grounds of quality control and safety of transplantable organs^46^. In this context, our study which deals with the preparation and characterization of blood vascular scaffold in an *in vivo* set up has immense potential which if carefully modulated can lead to development of implants for human in the clinics.

## Author contributions

**Debashish Banerjee:** Conceptualization, Methodology, Investigation, Writing - original draft **Nikhil B. Nayakawde:** Conceptualization, Methodology, Investigation. **Deepti Antony**, **Meghshree Deshmukh, Sudip Ghosh, Carina Silbohm and Evelin Berger:** Methodology, Investigation **Uzair Ul Haq:** Methodology and **Michael Olausson:** Methodology, Supervision, Funding acquisition.

## Acknowledgments

The authors would like to thank Rafael Camacho, Center for Cellular Imaging, University of Gothenburg for developing the image analysis pipelines, Dr. Anikó Kovács (A.K.), pathologist, Sahlgrenska University Hospital, University of Gothenburg for the Histological interpretation and Dr. Abdulhussain Haamid, University Veterinarian at Sahlgrenska Academy at Gothenburg University for animal welfare and supervision. We acknowledge the Centre for Cellular Imaging at the University of Gothenburg and the National Microscopy Infrastructure, NMI (VR-RFI 2019-00217) for providing assistance in microscopy.

## Data availability

1. The Mass spectrometry proteomics data have been deposited to the ProteomeXchange Consortium^47^ *via* the PRIDE (dataset identifier PXD021179).
2. The source code for all image analysis pipelines for Cell and Area work is available *via* github (https://github.com/CamachoDejay/BanerjeeD_cell_area_tools). This github repository is private (restricted access) until the acceptance of the manuscript, after which the repository will become publically available under the GNU General Public License v3.0. To grant reviewers access to the code, a separate file as a “.zip” folder containing an integrate copy of the source code can be provided during the time of review.

