## Supplementary file for "Differential fate of acellular vascular scaffolds in vivo involves macrophages and T-cell subsets"

###### 1(A) Animal handling and operation

During the current research the ARRIVE (Animals in Research: Reporting *In Vivo* Experiments)**^1^** guideline was adopted. For the compassionate use of animals AVMA guideline**^2^** and Grimace scale**^3^** were also followed during the entire duration of research to maintain the animal welfare and care as well as improve standards of reporting. Lewis and Norwegian Brown donor male, and Lewis female recipient rats (2-2.5 months’ age, weight ≈180-220 g, Janvier Labs, Saint Berthevin Cedex, France) were housed and bred with food and water *ad libitum* at the Gothenburg University Experimental Biomedicine Center as per approved Animal Ethical Guidelines (Göteborgs Djurförsöksetiska Nämnd, Ethical Number, 151/14) which is regulated and governed by Swedish law and regulations, as well as by EU directives on animal welfare. A one-week acclimatization period was given to the experimental rats to prevent stress-induced disease.

###### 1 (B) Preparation of biological scaffold

###### Harvesting of aorta –Surgical preparation

The donor male Lewis and Brown Norway rats (n=6-8) were operated under general anesthesia with inhalant anesthetic, Isoflurane. A non-steroidal anti-inflammatory agent was used as pre-operative analgesia (Carprofen @5 mg/kg SC). Subsequently, the rats were subjected to induction of anesthesia ina box (receptacle) with access to anesthesia vapor using a precision vaporizer (Isoflurane - 5% in 25% oxygen at a flow of 1 l/min) leading to loss of complete consciousness and pain. Loss of pain was monitored from the absence of any writhing-type movements and pain reflexes (toe pinching) and consciousness from the absence of the definite indicators of consciousness e.g. absence of head, body and eye reflexes as a response to touch **(Indicators of consciousness and pain**). The rats were then maintained under continuous isoflurane anesthesia (3%) using a facemask when a long midline incision was performed to expose the abdominal tract, lungs and the heart under dorsal recumbence. Diaphragm was cut open to expose heart and aorta. Collection of blood was immediately performed from the heart using a 1ml syringe and a 23G needle. The branches of aorta were ligated and the aorta was subsequently harvested. Animal was thus euthanized during the procedure of exsanguination performed under deep surgicalanaesthesia. The whole operative procedure did not take more than 15 minutes to complete.

###### 1 (C) Storage of harvested aorta

The harvested aorta/scaffold was rinsed in phosphate- buffer saline (PBS) containing antibiotic (0.5% penicillin, 0.5% streptomycin), and antimycotic (0.5% amphotericin B) finally washed with PBS and water before freezing them at -150^0^C within 4 h of death. Aortas were thawed overnight at 4°C and used either as donor control normal (untreated) and or used to produce donor control chemically decellularized aorta/scaffolds. The normal and decellularized scaffolds were further used for implantation as described below.

###### 1 (D) Sterilization of normal and chemically decellularized scaffolds

The normal or untreated donor untreated scaffolds after thawing was sterilized for an additional one cycle treatment with phosphate buffered saline (PBS) containing antibiotics (0.5% penicillin, 0.5% streptomycin) and antimycotic (0.5% amphotericin B) (Cat. No. A5955, Sigma) finally washedwith PBS and water to remove traces of antibiotics. The decellularized donor aortic scaffold was sterilized by a pH-adjusted (pH- 7-7.4) solution of peracetic acid (2%) and ethanol (20%)for 4 hours at room temperature followed by DNase treatment to achieve the recommended level of DNA load after decellularization.**^4^** The decellularized scaffolds were washed with PBS for 12 h. In total 3 cycles of treatment with peracetic acid-ethanol with an alternate cycle of PBS wash were given. The final PBS wash from both the decellularized and untreated aorta was collected aseptically and submitted to Clinical Microbiology Laboratory, Gothenburg University for detection of the presence of bacteria, mycoplasma, and fungus.

Upon confirmation of decellularization and sterilization, the chemically-treated aortas are designated as donor decellularized aorta (DLDC/DBNDC) in comparison to the donor normal untreated aorta (DLN and DBNN) in all subsequent sections. The two types of donor aortas are referred to as the control aorta. The recipient animal was prepared for surgery using the same **Indicators of consciousness and pain** as described for the preparation of donor aorta following the same analgesic (Carpropen, 5 mg/kg bwt, SC, 20-25 min preoperatively) and anesthetic (Isoflurane- 5% in 25% oxygen at a flow of 1 l/min) procedure for induction of analgesia and anaesthesia respectively. In addition, to prevent corneal drying and trauma, ophthalmic ointment (Viscotears) was applied in both eyes and recipient animals were provided with electric heating pads during the maintenance of anaesthesia and operative procedure. The animals were operated under general anaesthesia (Isoflurane- 3% in 25% oxygen at a flow of 1 l/min). An abdominal midline incision was used for implantation of sterile implant (1.5-2 cm in length) of normal and chemically decullarized DC aortic tissue.

To minimize the number of animals, two pieces each (≈ 2 cm) of either normal (DLN or DBNN) and decellularized donor (LDC or BNDC) scaffolds were implanted in the recipient animals. The 1^st^ piece was implanted in the omentum, and the second scaffold was placed subcutaneous (SC). Each implant was secured with 7/0 Prolene suture and when completed, the abdominal muscle was closed with continuous suturing followed by a closure of the skin using interrupted sutures (4-0) in two separate layers. The recipient rats during the whole operative procedure was constantly monitored to avoid excessive depression of cardiac and respiratory functions, or insufficient anesthesia from the chest and flank movements. Normal pink colour of the mucous membrane was also monitored. Body temperature was also monitored with a rectal thermometer. During recovery, each animal received 1-5 ml saline (SC). Animals were placed under a heating lamp and monitored until they are fully recovered or awake. Animals were allowed to recover on paper towels in a clean cage. The rats were provided continuously supplemental heat during recovery. When the animal attained ambulatory stage it was returned to original cage with immediate access to food and water. Softened food pellets for initial few days’ post-operation was also provided on cage floor.

The animals were monitored post-operative twice a day for three days with the administration of analgesics, Carpropen (5 mg/kg bwt, SC, once a day). Post-operative pain was carefully monitored by qualified staffs with knowledge of care and handling of laboratory animals. The level of spontaneous activity, back arching/ horizontal stretching, abdominal writhing, falling/staggering, poor gait and twitching, grooming, porphyrin secretions (ocular/nares), squint-eyed, pale eyes, pilo erection, teeth grinding, reduced food and water intake, polyphagia of bedding and increased aggressiveness if any when handled were monitored. Any decrease in body weight (more than 10%), and deviation from the behavioural, attitudinal and physiologic signs of pain were set as an exclusion criterion for termination or sacrifice of the animal. The recipient Lewis rats were sacrificed after two months of scaffold implantation in the same way as how donor surgery was performed. The implants from the subcutaneous and omentum were harvested prior to sacrifice and processed for further analysis as described under **Processing of Samples**. In addition to animal groups mentioned above, we also prepared a sham group consisting of female rats of same age as the recipient animals. The sham female Lewis rats (SHAM) were prepared for surgery similar to the recipient Lewis rats, but without implantation. Hence, similar surgical intervention that included the opening and closing the abdomen with the same anaesthetics and analgesics. These rats were sacrificed at the same time points as the other recipient groups and used for comparison of the serum cytokine status.

**1 (E) Indicators of consciousness and pain**

The level of consciousness was monitored from the 6 indicators of definite consciousness like (i) standing posture, (ii) head or body righting reflex, vocalization, (iv) eye blinking, (v) eye pursuit and (vi) response to external threat or menace (response to firm toe pinch)**^2^.** When there was absence of all these reflexes, operative procedure was initiated. In addition, the recipient rats’ post-operation and recovery were observed for any symptoms of pain according to the Rat Grimace Scale**^3^** which included absence of orbital tightening, no nose or cheek flattening, normal ear folds as well as whiskers in normal position. The entire surgical procedure was performed successfully and all the recipient as well as the sham treated animals survived at the end of 2 months of study.

###### 2. Immunohistochemical assessment

The formalin-fixed paraffin sections (4 μM) were deparaffinized subjected to heat mediated antigen retrieval for 25 min in 10 mM citrate buffer, pH 6. Sections were blocked with Candor Blocking solution (Cat. No. 110 050, CANDOR Bioscience GmbH, Wangen, Germany) +15% Normal Goat Serum (Code-005-000-121, Jackson ImmunoResearch Europe Ltd, Cambridge House, St. Thomas' Place, Ely CB7 4EX, UK) for 45 min, probed with the primary antibodies diluted in low cross buffer (**Supplementary Table S1)**, washed 4 times with PBS-Tween 20 (0.05%), and probed for 1h with secondary antibody, conjugated with a fluorescent probe (see **Supplementary Table S1**). Finally, the air-dried slides were mounted in fluorescent mounting media containing DAPI (Fluoroshield, Cat. No. ab104139; Abcam, Kingsfordweg 151, Amsterdam 1043GR, Netherlands) and imaged with a Epifluorescence Microscope (DM5100, Leica MicrosystemsTM, Wetzlar GmbH. Germany). The overlapped images at 20x magnification **(Supplementary** **Fig. S7-S10)** and for clarity and localization of cells 40x magnification (**Supplementary Fig. S11)** has been presented.

###### 3. Image acquisition and analysis

### The image analyses were performed using the pipelines developed by the Imaging Core Facility, Gothenburg University. The source code for all image analysis pipelines for Cell and Area work is also available at github (https://github.com/CamachoDejay/BanerjeeD_[cell_area_tools](https://github.com/CamachoDejay/BanerjeeD_cell_area_tools)). The image analysis involved the identification of (i) the area covered by target cells (immune cells) as well as the (ii) area stained by target ECM. During experimental acquisition, fluorescent images with 3 distinct configurations were taken named: Ch00 - nuclear staining, Ch01, and Ch02 depending on the immune markers or the ECM proteins and is specified in the corresponding figure legends. The overlapped images have been shown for clarity and localization of the area stained for different ECM components and macrophages.

###### (A) Cell work, MATLAB:

The cell analysis tool was designed to help assessing if the cells of an experimental condition were brighter than those of a control sample.

During experimental acquisition, fluorescent images are taken with 3 distinct configurations (colour channels): Ch00: nuclear staining, blue, DAPI; Ch01: red or green; Rhodamine RedX or AF488 (depending on the immune markers and specified in the corresponding figure legends), and Ch02 orange or green; Rhodamine RedX or AF488 (depending on the immune markers and specified in the corresponding figure legends. Due to the nature of the image quality obtained using epifluorescence microscopy, perfect nuclear segmentation was very challenging. Therefore, instead of implementing our image analysis pipeline *via* a cell by cell basis (which relies on accurate single-cell segmentation), we decided to base our calculations *via* a pixel wise strategy. During the image analysis, each channel was loaded and treated as follows.

###### Step 1: Determining the intensity levels expected for the control sample:

1) **Segmentation of the cell area using nuclear staining.** Ch00 is loaded. A median filter is implemented as a pre-processing method (9-pixel window). Segmentation is done viathe Otsu method.

###### 2) Recording the pixel wise intensity of Ch01 and Ch02 for all areas recognized as cells

(obtained in step 1.1). Consider this a long vector of numbers.

3) **Determining threshold values for the control condition.** This is done independently for Ch01 and Ch02 according to a user-defined confidence interval. For example, for a confidence interval of 0.99, then a threshold value is selected such that only 1% of theintensity values of the control are above it.
4) **Determining the random probability that both Ch01 and Ch02 are above threshold.** This is done by calculating in the control sample the probability that both Ch01 and Ch02 are above their respective thresholds.

###### Step 2: Determining if the intensity levels of the experiment are above those of the control:

1) **Segmentation of the cell area using nuclear staining.** Same method as in step 1.1.
2) **Recording the pixel wise intensity of Ch01 and Ch02 for all areas recognized ascells**. Same method as in step 1.2.
3) **Determining the percentage of pixels of Ch01 and Ch02 in the experiment that arebrighter than in control conditions.** This is simply done by calculating the percentage

of the total pixels recognized as cells that are above the threshold values calculated in step 1.3. A sample is considered to be brighter than control if its number of bright pixels is larger than that expected by the previously defined confidence interval.

4) **Determining the percentage of pixels that are simultaneously above the intensity threshold for Ch01 and Ch02.** A sample is considered to be simultaneously brighter on both Ch01 and Ch02 if the value determined in this step is above the random chance determined in step 1.4.

###### (B) Area work, MATLAB:

To assess (area distribution of a particular ECM component), we developed an image analysis pipeline to establish the areas of an experimental condition that were brighter than those expected by the control conditions. During experimental acquisition, images of a desired region are obtained for both the control and experimental conditions. These linked images are referred to as a data set. At each location, 3 distinct microscopy configurations where implemented: Ch00 – DAPI or nucleus staining, Ch01 –green, and Ch02 – red. While all 3 channels are loaded during analysis calculations are only done for a specific channel (user-defined). A data set is treated as follows:

###### Step 1: Determining the intensity levels expected for the control sample:

1) **Load and display fluorescent images for the control condition:** Ch00, Ch01, and Ch02.
2) **Calculate intensity descriptors for the desired channel.** The channel requested by the user is then extracted (Ch XX), and the mean and standard deviation of its intensity values are	calculated.
3) **Determining threshold values for the control condition (T).** This is done by calculating:

𝑇 = 𝑚𝑒𝑎𝑛 + 𝑐 x 𝑠𝑡𝑑

where, c is a user-defined scaling factor (e.g. 6)

###### Step 2: Quantify the area of the image which is brighter than control

1) **Segmenting bright areas in control and experimental condition.** As a segmentation strategy all pixels with intensity above the value of T are labelled as 1 and all other to 0.
2) **Quantifying the areas of control and experiment that are bright.** The segmented image is then used to calculate the areas of the image that are above the intensity threshold. The area is quantified in terms of pixels and square micrometer.

In some cases, we do not wish to treat the imaged area as a whole. Instead, we are interested in implementing our image analysis pipeline only at a specific region of interest (ROI). If this is the case then the user is required to specify the ROI at the beginning of the pipeline, in both control and experimental conditions.

###### 4. XY chromosome quantification- identifications of the resident cells around the implant

DNA was isolated from tissue samples (~10 mg) using the manufacturer’s protocol (DNeasy blood and tissue kit, Qiagen, Cat. No. 69504), quantified using NanoDrop One (Thermo Fisher), and analyzed for detection of X and Y chromosome using X and Y chromosome-specific primers**^5^.** ddPCR was performed using QX200™ Droplet Digital PCR System (Bio-Rad). EvaGreen assay was performed according to the manufacturer’s protocol using X (Fwd: CCCCCTCCCTGCAAGTTTT; Rev: TTCCACCGAATTCCCTTATCC) and Y (Fwd: AAGCCTTACAGAAGCCGAAAAA; Rev: TGTGGCACTTTAACCCTTCGA) chromosome specific primers (RxnReady, IDT, Inc. Coralville, Iowa, USA). Each 20 μl reaction contained 10 μl 2x ddPCR supermix for EvaGreen (Cat. No. 186-4034, Bio-Rad), 1 μl of either X or Y chromosome specific primer and 2 μL of gDNA template. Droplets were generated on the QX100 droplet generator (Bio-Rad, cat. No. 186-4008,) using droplet generation oil for EvaGreen (Bio-Rad Cat. No., 1864005), transferred to 96-well plates (Bio-Rad) and sealed with a pierce-able foil heat seal (Bio-Rad). PCR was performed in C1000 Touch™ thermal cycler (Bio-Rad) under the following cycling conditions: 1× (95°C for 5 min), 40× (95°C for 30s, 60°C for 1 min), 1× (4°C for 5 min, 90°C for 5 min), 1× (4°C until the plate is read) with a ramp rate of 2°C/s. The plate was incubated overnight at 4°C prior to being read using a QX200 droplet reader (Bio-Rad). The percentage of Y chromosomes were calculated from the X and Y chromosome copy number as -

Y (%) = (Y chromosome copy number) / (X + Y chromosomes copy number) x 100

**5. RNA Quantification and ddPCR**

The untreated and implants were thawed, weighed, lyophilized, and cut into about 5 mg pieces and subjected to homogenization (4,0 m/s for 40 s, 2 cycles; FastPrep^TM^ system-24 5G, MP Biomedicals, CA). RNA was isolated using RNeasy®micro kit (Cat No. 74004, Qiagen Str. 1, 40724 Hilden, Germany), measured usingNanoDrop One (Thermo Fisher Scientific, 168, MA USA 02451), integrity assessed (Agilent RNA 6000 Pico Kit, Agilent 2100 Bioanalyzer instrument), and used finally for cDNA conversion (Cat No. 1708891; iScriptTM cDNA synthesis kit; BioRad, Alfred Nobel Drive Hercules, California 94547, USA) as per the manufacturer’s instructions.

A ddPCR targeting CD68 (dRnoCPE5149097, Bio-Rad), CD163 (dRnoCPE5175377, Bio- Rad), MRC1 (dRnoCPE5170185, Bio-Rad), IL-10 (dRnoCPE5172671, Bio-Rad) and beta Actin (β-actin) (Actb, Rn.PT.39a.22214838.g, Integrated DNA Technologies (IDT), Inc. Coralville, Iowa, USA) as the reference gene was performed using QX200™ Droplet Digital PCR System (Bio-Rad) and TaqMan based hydrolysis probe.**^6^** Each of the 20 µl reactions contained 1×ddPCR supermix for probes (Cat. No. 1863024, BioRad), 1 µl of HEX primer- probe mix, and 2 µl of cDNA. Reaction mixtures were prepared in a semi-skirted 96- well plate. Droplets were generated on the QX100 droplet generator (Bio-Rad, cat. No. 186-4008,) using droplet generation oil for probes (Bio-Rad, Cat. No. 1863005), transferred to 96-well plates (Bio-Rad) and sealed with a pierce-able foil heat seal (Bio-Rad). PCR was performed in a C1000 Touch™ thermal cycler (Bio-Rad). The plate was then incubated at 4°C overnight prior to being read using a QX200 droplet reader (Bio-Rad). Every ddPCR run included negative template controls (NTCs) in duplicates.

QuantaSoft™ Analysis Pro software (version 1.0.596) was used for data analysis. Thresholds were manually set for each sample using acceptance criteria previously defined. The target gene expression value was represented with modification as copies/ng of calculated input RNA**^7^** (**Supplementary fig. S10).**

###### 6. Proteomic Analysis

For proteome analysis, 3-4 mm of donor scaffolds and implants were used. In total, twelve groups were selected for proteomic analysis **(Supplementary Table S2)**.

###### (A) Sample Preparation

The samples were homogenized in the lysis buffer (50 mM triethylammonium bicarbonate (TEAB), 2% sodium dodecyl sulfate (SDS), centrifuged, and supernatants transferred into new vials. Protein concentration was determined using Pierce™ BCA Protein Assay (Cat. No. 23225, Thermo Fisher Scientific, 168, MA USA 02451). As reference samples, the respective normal controls were chosen to be analyzed in both TMT sets. Aliquots containing 30 μg of each sample and the references were digested with trypsin using the filter-aided sample preparation (FASP) method**^8^**. Peptides were collected by centrifugation. Digested peptides were labeled using TMT 11-plex isobaric masstagging reagents (Thermo Fisher Scientific, 168, MA USA 02451) according to the manufacturer's instructions. The combined samples of each of the two TMT sets were pre- fractionated with basic reversed-phase chromatography (bRP-LC) using a Dionex Ultimate 3000 UPLC system (Thermo Fisher Scientific, 168, MA USA 02451). Peptide separations were performed using a reversed-phase XBridge BEH C18 column (3.5 μm, 3.0x150 mm, Waters Corporation) and a linear gradient from 3% to 40% solvent B over 17 min followed by an increase to 100% B over 5 min. Solvent A was 10 mM ammonium formate buffer at pH 10.00and solvent B was 90% acetonitrile, 10% 10 mM ammonium formate at pH 10.00. The 40 fractions were concatenated into 20 fractions, dried, and reconstituted in 3% acetonitrile, 0.2% formic acid.

###### (B) Liquid Chromatography-Tandem Mass Spectrometry Analysis

The fractions were analyzed on a QExactive HF mass spectrometer interfaced with the Easy- nLC^TM^ 1200 liquid chromatography system (Thermo Fisher Scientific, 168, MA USA 02451).

Peptides were trapped on an Acclaim Pepmap 100 C18 trap column (100 μm x 2 cm, particle size 5 μm, Thermo Fisher Scientific, 168, MA USA 02451) and separated on an in-house packed analytical column (75 μm x 300 mm, particle size 3 μm, Reprosil-Pur C18, Dr. Maisch) using a gradient from 7% to 35% B over 76 min followed by an increase to 100% B for 8 min at a flow of 300 nL/min. The instrument operated in a data-dependent mode where the precursor ion mass spectra were acquired at a resolution of 60 000, the 10 most intense ions were isolated in a 0.7 Da isolation window and fragmented using collision energy HCD settings at either 28 or 50. MS2 spectra were recorded at a resolution of 60 000, charge states 2 to 4 were selected for fragmentation and dynamic exclusion was set to 15 s with 10 ppm tolerance.

###### (C) Protein Identification and Quantification

The data files for each TMT set were merged for identification and relative quantification using Proteome Discoverer^™^ version 2.2 (Thermo Fisher Scientific, 168, MA USA 02451). The database search was performed using the Swissprot Rat Database (Oct 2018) using Mascot 2.5.1 (Matrix Science) as a search engine with a precursor mass tolerance of 5 ppm and fragment mass tolerance of 200 mmu. Tryptic peptides were accepted with zero missed cleavage, variable modifications of methionine oxidation and fixed cysteine alkylation, TMT- label modifications of N-terminal and lysine were selected. Percolator was used for the validation of identified proteins and the quantified proteins were filtered at 1% FDR and grouped by sharing the same sequences to minimize redundancy. Reporter ion intensities were quantified in MS2 spectra using the S/N values as abundances and normalized on the total protein abundance within the Proteome Discoverer 2.2 workflow. Only peptides unique for a given protein were considered for quantification and ratios were calculated by normalizing the samples with the reference normal control samples. Only proteins quantified in all samples and relative standard deviation less than 25 percent in either syngeneic or allogeneic implant datasets were considered for comparisons among different sample groups. Proteins with two-fold expression changes were selected as up- or down-regulated.

###### (D) PPI network construction and module analysis

Employing the Search Tool for the Retrieval of Interacting Genes (STRING v.10.5)**^9^,** we built the PPI network of the identified proteins. The minimum required interaction score was set to 0.400 (medium confidence), and disconnected nodes (proteins with no known interactions) were not included in the view. Moreover, the MCODE plugin of Cytoscape (v.3.7.2)**^10^** was used to identify network modules with degree cut-off = 2, node score cut- off = 0.2, k-core = 2, and max. depth = 100. The topological features of the modules were determined *via* NetworkAnalyzer**^11^** and Cytohubba**^12^** plugins of Cytoscape to identify the hub proteins. The hub proteins were mapped to the ECM and Immune responsive network modules. Pathway enrichment analyses of the networks or modules were performed through DAVID (v.6.8) **^13^** using KEGG**^14^.** Benjamini-Hochberg’s adjusted p-values <0.05 were considered for statistically significant enrichment results. Gene Ontology (GO) enrichment analyses were also performed for terms related to the extracellular matrix (ECM) and immune responses.

###### (E) Proteome Profiler of cytokines present in serum

Briefly, the Proteome Profiler Rat Cytokine Array Kit, column A contains 4 membranes coated with 29 cytokines/chemokines in duplicate spots. 150 μl of serum from the recipients and SHAM rats was mixed with a cocktail of biotinylated detection antibodies and compared separately. The nitrocellulose membranes were subsequently blocked according to the manufacturer's protocol. The protein and antibody mixtures were then incubated with the membrane containing immobilized antibodies for 29 rat cytokines. Bound protein was detected with streptavidin conjugated to horseradish peroxidase (HRP). Membranes were washed and developed with chemiluminescent detection reagents. The dot blot membranes were analyzed for the cytokine/chemokine spots generated as chemiluminescent signals which were detected using Azure C600 instrumentation. The chemiluminescent signal was expressed as Mean pixel intensity using Fiji as an image processing package**^15^.** For each membrane, the chemiluminescent signals generated from each of two paired spots (representing a single cytokine) were normalized to negative and positive reference spots as elaborated**^16^.**

**7. Statistical analysis**

Samples from different groups (non- implanted vs implanted) were subjected to parametric and non-parametric statistics after detection of normal distribution by visual observation following calculation of descriptive statistics, Q-Q plots, and frequency distribution from the Clean Data as described below. In addition, the data were subjected to the Shapiro-Wilk test for normality distribution to accept or reject the null hypothesis that the sample has been drawn from a normally distributed population. Outliers, which may increase the variability in the non-ordinal data was removed by using the ROUT method (Q set to 1%). Cleaned data was used to find normal distribution.

When the population under study consisted of all normally distributed groups, a One-way ANOVA was performed to compare 3 or more groups, with Dunnett's T3 Multiple Comparisons post hoc test. Results are presented as Mean±SEM. When the population consisted of ordinal data (histology scores) and non-normal population distribution, the Kruskal-Wallis test was performed to find out the statistical difference between the groups with Dunn's Multiple Comparisons post hoc tests. Results are presented as the Max and Minimum value with the median value being shown as a horizontal line at the centre of individual box- whisker. The statistical analysis carried out for each experiment is defined in the corresponding figure legends.
